# Plasmid-mediated colistin resistance among human clinical *Enterobacterales* isolates: National surveillance in the Czech Republic

**DOI:** 10.1101/2023.02.09.527831

**Authors:** Marketa Zelendova, Costas C. Papagiannitsis, Petra Sismova, Matej Medvecky, Katarina Pomorska, Jana Palkovicova, Kristina Nesporova, Vladislav Jakubu, Ivana Jamborova, Helena Zemlickova, Monika Dolejska, Working Group for Monitoring of Antibiotic Resistance

## Abstract

The occurrence of colistin resistance has increased rapidly among *Enterobacterales* around the world. We performed a national survey of plasmid-mediated colistin resistance in human clinical isolates by a retrospective analysis of samples from 2009-2017 and a prospective sampling in 2018-2020. The aim of this study was to identify and characterize isolates with *mcr* genes from various regions of the Czech Republic using whole genome sequencing (WGS). Of all 1932 colistin-resistant isolates analyzed, 73 (3.8%) were positive for *mcr* genes. Most isolates carried *mcr-1* (48/73) and were identified as *Escherichia coli* (n=44) and *Klebsiella pneumoniae* (n=4) of various sequence types (ST). Twenty-five isolates including *Enterobacter* spp. (n=24) and *Citrobacter freundii* (n=1) carrying *mcr-9* gene were detected, three of them (*Enterobacter kobei* ST54) co-harbored the *mcr-4* and *mcr-9* genes. Multi-drug resistance phenotype was a common feature of *mcr* isolates and 14% (10/73) isolates also co-harboured clinically important beta-lactamases including 2 isolates with carbapenemases KPC-2 and OXA-48. Phylogenetic analysis of *E. coli* ST744, the dominant genotype in this study, with the global collection, showed Czech isolates belonged to two major clades, one containing isolates from Europe, while the second composed of isolates from diverse geographical areas. The *mcr-1* gene was carried by IncX4 (34/73, 47%), IncHI2/ST4 (6/73, 8%) and IncI2 (8/73, 11%) plasmid groups. Small plasmids belonging to the ColE10 group were associated with *mcr-4* in three isolates while *mcr-9* was carried by IncHI2/ST1 plasmids (4/73, 5%) or the chromosome (18/73, 25%). We showed an overall low level of occurrence of *mcr* genes in colistin-resistant bacteria from human clinical samples in the Czech Republic.

## Introduction

The excessive consumption of antimicrobial substances associated with faster spread of antibiotic resistance represents a global concern. The dissemination of multi-drug resistant (MDR) bacteria resulted in limited treatment options of infectious diseases in healthcare systems. The interest in the administration of older antibiotics such as polymyxins has been therefore renewed (Bitar et al., 2020). Colistin has been widely used in the past, but due to its nephrotoxicity and neurotoxicity it has become a restricted antibiotic (Vińes et al., 2021). Currently, colistin is administered for the treatment of life-threatening infections caused by MDR Gram-negative pathogens as the last-resort antibiotic (Yilmaz et al., 2016; Hamel et al., 2021; Vińes et al., 2021). In contrast, colistin has been widely used for prophylactic and therapeutic purposes in veterinary medicine for decades (Quiroga et al., 2019; Vińes et al., 2021). However, colistin overuse in livestock has led to the spread of colistin-resistant pathogens worldwide and the development of different strategies used by bacteria to increase resistance against colistin (El-Sayed Ahmed et al., 2020).

Resistance to colistin can be either associated with chromosomal mutations or with *mcr* genes carried by plasmids that are facilitating horizontal transfer of colistin resistance between bacteria (Zhu et al., 2019). Acquired colistin-resistance mechanisms have been recognized in some members of Enterobacteriaceae family, such as *E. coli, Salmonella* spp., *Klebsiella* spp., and *Enterobacter* spp. These include genes and operons responsible for encoding enzymes that have a direct role in LPS modification, such as the *pmrC* and *pmrE* genes and the *pmrHFIJKLM* operon (Aghapour et al., 2019). Apart from the chromosomally-mediated mechanisms, 10 variants, *mcr-1* to *mcr-10*, carried by various plasmid families have been so far identified in *Enterobacterales*, especially in *E. coli* and *Enterobacter* spp. (Bitar et al., 2020; Li et al., 2020; Javed et al., 2020; Wang et al., 2020). The most common variant, *mcr-1*, is usually located on plasmids of various incompatibility (Inc) groups, but predominantly on IncX4, IncI2 and IncHI2 (Caratolli et al., 2014; Doumith et al., 2016; Zelendova et al., 2021). These plasmid types carrying *mcr-1* were found in *Enterobacterales* isolates from humans as well as farm animals around the globe (Dalmolin et al., 2018; Quoriga et al., 2019), highlighting their wide distribution in various niches. Besides *mcr-1*, other genes for plasmid-mediated colistin resistance, such as *mcr-4* and *mcr-9*, have been reported (Bitar et al., 2020; Li et al., 2020). The *mcr-4* gene is usually located on small ColE10-type plasmids (Caratolli et al., 2017; Marchetti et al., 2021) while *mcr-9* is mostly carried by large IncHI2 plasmids or is incorporated into the chromosome (Li at al. 2020; Tyson et al., 2020).

The emergence of colistin resistance in MDR bacteria is a significant clinical concern. Isolates encoding extended-spectrum beta-lactamase (ESBL) or carbapenemase on a single plasmid along with *mcr* have been detected (Caratolli et al., 2014; Katip et al., 2021). As the co-occurrence of more resistance genes within the bacteria represents a threat for current medicine, The European Centre for Disease Prevention and Control (ECDC) published the expert protocol that recommends to perform the surveillance of co-resistance to both colistin and carbapenems in *Enterobacterales* (ECDC technical report).

From the Czech Republic, only sporadic reports describing the identification of *mcr*-carrying isolates in clinical samples have been published so far (Bitar et al., 2019; Bitar et al., 2020; Krutova et al., 2021), however, overview data on prevalence of *mcr* genes in Czech patients are not available. To fill in this gap, we aim to identify *mcr* genes in colistin-resistant human clinical isolates of Gram-negative bacteria from the Czech Republic between 2009 and 2020, and to determine characteristics of the *mcr*-positive strains using whole genome sequencing (WGS), plasmid typing and transferability experiments.

## MATERIALS AND METHODS

### Sampling and detection of *mcr* genes

A total of 1932 colistin-resistant isolates of Gram-negative bacteria with minimum inhibitory concentration (MIC) to colistin >2 mg/L collected from Czech patients between 2009 and 2020 were examined. The collection consisted of 682 retrospective isolates obtained from 2009 till 2017 during various surveillance programs at the National Institute of Public Health that were not targeting colistin resistance. The retrospective collection included mainly *Klebsiella pneumoniae* (n=429), *Enterobacter* spp. (n=108), *Pseudomonas aeruginosa* (n=49), *Acinetobacter baumanii* (n=33), *Stenotrophomonas maltophilia* (n=20), *Escherichia coli* (n=14) and 29 isolates of other 13 species. Prospective surveillance of colistin resistance in Czech hospitals was carried out during 2,5-year period between January 2018 and June 2020. It resulted in the collection of 1250 isolates of *Klebsiella pneumoniae* (n=491), *Enterobacter* spp. (n=311), *Escherichia coli* (n=179), *Pseudomonas aeruginosa* (n=99), *Acinetobacter baumanii* (n=43), *Hafnia alvei* (n=28), *Klebsiella variicola* (n=20), *Acinetobacter* spp. (n=15), *Salmonella enterica* (n=15), *Klebsiella oxytoca* (n=14), *Klebsiella aerogenes* (n=10), and 25 isolates of other 11 species. Strain identification was performed by matrix-assisted laser desorption ionization-time of flight mass spectrometer (MALDI-TOF) using MALDI Biotyper software (Bruker Daltonics, Bremen, Germany). All isolates were subjected to multiplex polymerase chain reaction (PCR) to detect the variant of *mcr* genes (*mcr-1* to *mcr-9*) (Rebelo et al., 2018; Kieffer et al., 2019; Wang et al., 2018).

### Antimicrobial susceptibility testing

Susceptibility profiles of *mcr*-positive isolates was determined by broth microdilution method using the following 15 antimicrobial substances: amikacin, ampicillin, ampicillin/sulbactam, cefepime, cefotaxime, cefoxitin, ceftazidime, ceftolozane/tazobactam, colistin, cotrimoxazole, ciprofloxacin, gentamicin, meropenem, piperacillin/tazobactam and tobramycin. The production of ESBL and AmpC type beta-lactamase was tested by double-disk synergy test (EUCAST 2017). The production of carbapenemase was tested by combination disc test method (EUCAST 2017) and biochemical tests (BioRad-Beta-Carba test) while carbapenem hydrolysis was tested by MALDI-TOF (Papagiannitsis et al., 2015).

### Conjugative transfer of *mcr* genes

Conjugation assays were performed to determine the transferability of *mcr* genes into plasmid-free sodium azide-resistant *E. coli* J53 K12 recipient cells using filter-mating method (Borowiak et al., 2019). The transconjugants (TCs) were selected on LB agar plates (LBA) with sodium azide (100 mg/L) and colistin (0.5 mg/L). Successful transfer of the plasmid-mediated colistin resistance via conjugation was confirmed by PCR targeting the *mcr* gene (Rebelo at al. 2018, Kieffer et al., 2019; Wang et al., 2018) and *E. coli* J53 K12 (Bauer et al., 2007). The size and number of plasmids transferred were estimated by pulsed-field gel electrophoresis (PFGE) using S1 nuclease (CDC 2004) and PCR-based replicon typing (PBRT; Carattoli et al., 2005).

### Whole genome sequencing and plasmid characterization

Genomic DNA of all *mcr*-positive isolates was extracted using NucleoSpin® Tissue kit (Macherey-Nagel GmbH & Co, Duren, Germany). The libraries were prepared using Nextera XT DNA Sample Preparation Kit and sequenced on MiSeq or NovaSeq 6000 platform (Illumina, San Diego, CA, USA). Raw reads were quality- and adaptor-trimmed using Trimmomatic v0.39 (Bolger at al., 2014) and assembly was performed by SPAdes v3.12.0 (Bankevich et al., 2012) and assembled data were analyzed using the CGE tools (https://cge.cbs.dtu.dk/) that were used to identify antibiotic resistance genes (ResFinder 4.1) (Zankari et al., 2012), multi-locus sequence types (MLST 2.0) (Larsen et al., 2012), plasmid replicons (PlasmidFinder 2.1) and plasmid sequence types (STs) (pMLST 2.0) (Carattoli et al., 2014). Chromosomal mutations for resistance to fluoroquinolones and colistin in *E. coli* and *K. pneumoniae* isolates were determined by PointFinder (Zankari et al., 2017). Sequences of six IncX4 plasmids carrying *mcr-1* were extracted from Illumina data and gaps were filled by PCR-based strategy and Sanger sequencing.

Complete nucleotide sequence of 12 selected isolates was obtained using long-read sequencing on MinION platform (Oxford Nanopore technologies, ONT, Oxford, UK). Genomic DNA was extracted by Genfind V3 (Beckman Coulter, USA). Libraries were constructed using a SQK-RBK004 rapid barcoding 1D kit according to the manufacturer’s protocol. The barcoded library mix was loaded onto a flow cell (FLO-MIN106 R9.4 SpotON) and sequenced for 48 h. The raw electrical signals were basecalled using Guppy v4.2.2 (ONT) and raw reads in fastq format were obtained. BBDuk (https://jgi.doe.gov/data-and-tools/software-tools/bbtools/bb-tools-user-guide/) and Porechop v0.2.4 (ONT) were used for adaptor and quality trimming (Q ≤ 7) and for demultiplexing, respectively. Whole plasmid sequences were assembled using Unicycler v0.4.8 (Wick et al., 2017) and Flye v2.6 (Lin et al., 2016) and polished by Illumina reads using Pilon v1.23 (Walker et al., 2014). For sequence analysis and annotation, BLAST (www.ncbi.nlm.nih.gov/BLAST), the ISfinder database, and the open reading frame (ORF) finder tool (www.bioinformatics.org/sms/) were used. Comparative genome alignment with corresponding reference plasmids was performed using Mauve v.2.3.1 (Darling et al., 2010). Figures were generated from sequence data using BRIG v.0.95 (Alikhan et al., 2011) and clinker v0.0.23 (Gilchrist and Chooi, 2021).

### Phylogenetic analysis

In total, four different datasets were subjected to phylogenetic analysis. Two of them were local phylogenetic trees including only isolates from our collection: the first one comprised all detected *mcr*-carrying *E. coli* isolates and the second one showed the phylogeny of *Enterobacter* spp. genomes. The third tree was global and comprised genomes of 449 *E. coli* ST744 isolates that were available at EnteroBase in April 2021 (http://enterobase.warwick.ac.uk/) along with ten ST744 isolates from our collection. These three trees were generated based on a core-genome determined employing a Roary pipeline v3.12.0 (Page et al., 2015) and aligned with MAFFT v7.313 (Katoch et al., 2013). Trees were inferred under GTR+CAT model using FastTree v2.1.11 (Price et al., 2010) compiled with double precision arithmetic.

Remaining detailed tree topology was constructed based on a pipeline described in previous study (Forde et al., 2022) using Python scripts that are available on GitHub (https://github.com/matejmedvecky/anthraxdiversityscripts). Based on *E. coli* ST744 global tree, 38 Illumina SRA archives belonging to isolates that were closely related to ten ST744 isolates from our collection were gathered from the GenBank database in May 2021. Raw sequencing reads of those 38 isolates along with another ten from our collection were subjected to quality trimming via Trimmomatic tool v0.36 (Bolger et al., 2014) and consequently mapped to *E. coli* str. K-12 substr. MG1655 reference genome (GenBank accession U00096.3) using Bowtie2 v2.3.4.2 (Langmead et al., 2012). Single nucleotide polymorphisms (SNPs) were detected in individual isolates by VarScan v2.4.4 (Koboldt et al., 2012) using following parameters: minimum read depth of 8; minimum base quality of 20; variant allele frequency ≥ 0.80. Problematic sites were then removed based on the following rules: occurred in phage regions as detected by PHASTER (Arndt et al., 2016); occurred in repetitive/homologous genomic regions; more than 5 isolates at a particular site showed prevalent base frequency below 80% or/and read depth below 8. Resulting alignment file was then subjected to maximum-likelihood analysis using RAxML v8.2.11 (Stamatakis, 2014) under GTR+CAT model of nucleotide substitution with 500 rapid bootstrap replicates using sample SRR9990292 as an outgroup. Tree topologies were visualised via iTOL v6.3 (Letunic and Bork 2021) and edited using Inkscape v1.1 (https://inkscape.org/cs/).

Species level discrimination of *Enterobacter* spp. was performed using average nucleotide identity (ANI) (Yoon et al., 2017) and digital DNA-DNA hybridization (Meier-Kolthoff et al., 2013) of whole genome sequences. Eight type strains were used as reference species including *Enterobacter asburiae* (ATCC35953^T^), *Enterobacter bugandensis* (DSM 29888^T^), *Enterobacter cloacae* ATCC13047^T^), *Enterobacter dykesii* (DSM111347^T^), *Enterobacter hormaechei* (ATCC49162^T^), *Enterobacter kobei* (DSM13645^T^), *Enterobacter vonholyi* (DSM110788^TT^), *Enterobacter roggenkampii* (DSM16690^T^).

### Nucleotide sequence accession numbers

Genome assemblies, SRA archives and annotated plasmid sequences (Table 1) were deposited in NCBI under BioProject with accession number PRJNA772899.

**Table 1.** The characteristics of sequenced *mcr*-encoding plasmids

| Plasmid name | Organism (ST)<sup>1</sup> | Plasmid-carrying *mcr*<sup>2</sup> | *mcr* | Other ARGs in *mcr* plasmid<sup>3</sup> | WGS platform | Accession no. |
| --- | --- | --- | --- | --- | --- | --- |
| pMCR1-40331 | *K. pneumoniae* (ST290) | IncX4 (33,303) | *mcr-1.1* | - | Illumina | OP428973 |
| pMCR1-42913 | *E. coli* (ST448) | IncX4 (33,304) | *mcr-1.1* | - | Illumina | OP428974 |
| pMCR1-44158 | *E. coli* (ST156) | IncX4 (33,303) | *mcr-1.2* | - | Illumina | OP428975 |
| pMCR1-44653 | *E. coli* (ST1196) | IncX4 (33,303) | *mcr-1.1* | - | Illumina | OP428976 |
| pMCR1-45082 | *E. coli* (ST744) | IncX4 (34,068) | *mcr-1.1* | - | Illumina | OP428977 |
| pMCR1-46049 | *K. pneumoniae* (ST147) | IncX4 (33,303) | *mcr-1.1* | - | Illumina | OP428978 |
| pMCR1-53288 | *E. coli* (ST538) | IncI2 (60,733) | *mcr-1.1* | - | MinION | OP434482 |
| pMCR1-43934 | *E. coli* (ST8186) | IncHI2/ST4 (225,732) | *mcr-1.1* | *aph(6)-Id, aph(3'')-Ib, catA1, tet(A)* | MinION | OP950834 |
| pMCR1-51133 | *E. coli* (ST117) | IncHI2/ST4 (237,743) | *mcr-1.1* | *aadA1, aadA2b, bla<sub>TEM-1B</sub>, catA1, cmlA1, qacE, sul1, tet(A)* | MinION | OP950835 |
| pMCR1-59496 | *K. pneumoniae* (ST726) | IncHI2/ST3 (254,909) | *mcr-1.1* | *aac(3)-IV, aadA1, aadA2b aph(3')-Ia, aph(4)-Ia, bla<sub>CTX-M-14</sub>, cmlA1, floR, fosA3, mph(A), sul2, sul3* | MinION | OP950836 |
| pMCR9-16539 | *E. kobei* (ST591) | IncHI2/ST1 (285,283) | *mcr-9.1* | *aadA2b, aph(3'')-Ib, aph(6)-Id, bla<sub>TEM-1B</sub>, bla<sub>SHV-12</sub>, catA2, dfrA19, qacE, qnrA1, sul1, tet(D)* | MinION | OP950838 |
| pMCR9-17620 | *E. hormaechei* (ST91) | IncHI2/ST1 (276,870) | *mcr-9.1* | *aadA2b, ant(2'')-Ia, bla<sub>CTX-M-9</sub>, catA1, dfrA16, qacE, qnrA1, sul1* | MinION | OP950833 |
| pMCR9-57185 | *C. freundii* (ST18) | IncHI2/ST1 (330,692) | *mcr-9.1* | *aac(6')-IIc, bla<sub>TEM-1B</sub>, bla<sub>DHA-1</sub>, catA2, ere(A), qacE, qnrB4, sul1, tet(D)* | MinION | OP950837 |
| pMCR4-26153 | *E. kobei* (ST54) | ColE10 (12,808) | *mcr-4.2* | - | MinION | OP428979 |

<sup>1</sup>ST, sequence type.
<sup>2</sup>Plasmid carrying *mcr* include the information on plasmid incompatibility group (Inc), ST (if available) and plasmid size in bp.
<sup>3</sup>ARGs, antibiotic resistance genes

## Results

### *mcr-*positive *Enterobacterales* isolates

From all 1932 examined colistin-resistant isolates, 73 (3.8%) were identified to carry *mcr* genes (Supplementary Table S1). Most (65/73) isolates were detected during the prospective years including eight isolates in 2018 (3%, n=274), 27 isolates in 2019 (4%, n=634) and 30 isolates in 2020 (9%, n=342). From the retrospective analysis using a collection of isolates at National reference laboratory for antibiotics, seven isolates (1%, n=682) carrying *mcr* genes were found including seven *Enterobacter* spp. from 2010 (n=1), 2012 (n=3), 2013 (n=1), 2014 (n=2) and one isolate of *K. pneumoniae* (n=1) from 2017. Isolates carrying *mcr-1* were identified as *E. coli* (n=44) and *K. pneumoniae* (n=4), while the remaining 25 isolates carried the *mcr-9*.*1* allele. Three of the *mcr-9*.*1*-carrying isolates were also positive for *mcr-4*.*2*/*mcr-4*.*3*. The isolates carrying *mcr-9*.*1* were identified as *Citrobacter freundii* (n=1), *Enterobacter asburiae* (n=13), *Enterobacter kobei* (n=6), *Enterobacter cloacae* (n=3). *Enterobacter roggenkampii* (n=1) and *Enterobacter hormaechei* (n=1). Plasmid-mediated colistin resistance genes were the most common among *E. coli* as 19% (44/231) colistin-resistant isolates carried *mcr-1* while the occurrence in other species was rare (4.4% in *Enterobacter* spp., 0.4% in *K. pneumoniae*).

Colistin-resistant isolates carrying *mcr-1* (48/73, 66%) showed phenotypic resistance to beta-lactam antibiotics including ampicillin (46/48, 96%), ampicillin/sulbactam (43/48, 90%), cefoxitin (9/48, 19%), piperacillin/tazobactam (9/48, 19%), cefotaxime (5/48, 10%), ceftazidime (5/48, 10%), cefepime (4/48, 8%) and ceftolozane/tazobactam (2/48, 4%). Resistance to other antimicrobials including cotrimoxazole (34/48, 71%), ciprofloxacin (34/48, 71%), trimethoprim (34/48, 71%), gentamicin (11/48, 23%), tobramycin (10/48, 21%) and amikacin (1/48, 2%) was found.

The majority of colistin-resistant isolates carrying *mcr-9* were resistant to cefoxitin (25/25, 100%), ampicillin (23/25, 92%), ampicillin/sulbactam (23/25, 92%) and cefotaxime (12/25, 48%). Furthermore, they showed resistance to ceftazidime (9/25, 36%), cotrimoxazole (6/25, 24%), ciprofloxacin (5/25, 20%), tobramycin (5/25, 20%), trimethoprim (5/25, 20%), piperacillin/tazobactam (7/25, 28%), gentamicin (4/25, 16%), cefepime (3/25, 12%), amikacin (2/25, 8%), ceftolozane/tazobactam (2/25, 8%) and meropenem (2/25, 8%). Nine isolates including *E. coli* (n=4), *Enterobacter* spp. (n=3), *Citrobacter freundii* (n=1) and *K. pneumoniae* (n=1) with resistance to seven or more different antibiotics were simultaneously positive for ESBL production. AmpC beta-lactamase was detected in six *Enterobacter* spp. isolates and one *E. coli* isolate.

### Analysis of WGS results

The *mcr-1*-positive *E. coli* isolates belonged to 26 various STs of which the *E. coli* ST744 was the most common (10/44). Fifteen *E. coli* isolates were assigned to ST88 (n=3), ST538 (n=3), ST1011 (n=3), ST69 (n=2), ST162 (n=2), and ST453 (n=2), while the remaining nineteen isolates belonged to unique STs (Table S1, Figure 1). Four *K. pneumoniae* isolates carrying *mcr-1* were assigned to four different STs (ST147, ST231, ST290 and ST726) and one *C. freundii* isolate with *mcr-9* belonged to ST18. *Enterobacter* spp. with *mcr-9* belonged predominantly to *E. asburiae* of two different STs including ST484 (n=11) and ST358 (2) (Figure 2). Six isolates of *E. kobei* belonged to ST32 (n=1), novel ST (n=1), ST591 (n=1) and ST54 (n=3). All *E. kobei* ST54 isolates originated from patients in a single hospital and carried *mcr-4* apart from *mcr-9*. Remaining two isolates were identified as *E. cloacae* ST1525 and were obtained from one patient.

**Figure 1.**
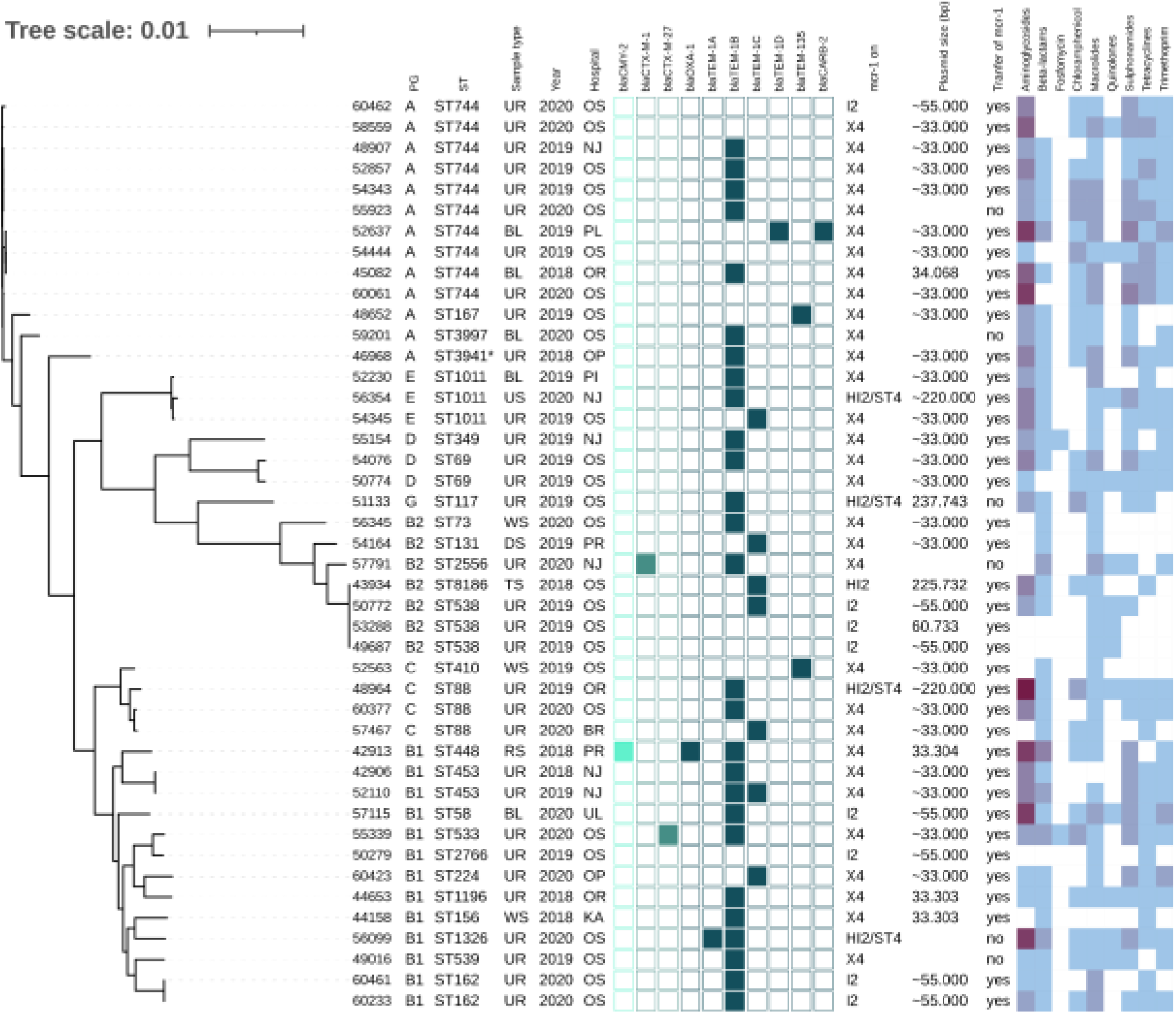
Phylogenetic tree of *E. coli* isolates with *mcr-1* of Czech clinical origin. The metadata in columns represents phylogenetic group (PG); sequence type (ST); type of sample (Sample type): urine (UR), blood (BL), rectal swab (RS), tonsil swab (TS), wound swab (WS), decubitus swab (DS), urethra swab (US); year of isolation (Year) and city where is the hospital related to the isolate recovery (Hospital): Novy Jicin (NJ), Prague (PR), Ostrava (OS), Karvina (KA), Ostrava-Poruba (OR), Opava (OP), Pribram (PI), Plzen (PL), Usti nad Labem (UL), Brno (BR). The turquoise squares represent presence (full square) or absence (empty square) or respective beta-lactamase encoding genes divided as AmpC (bright turquoise), ESBL (medium) and narrow-spectrum beta-lactamases (dark).The next section (mcr-1 on) reveals which plasmid carried *mcr-1* gene; the size of the plasmid (Plasmid size) in bp while approx. sizes (∼) are estimated based by S1-PFGE while the more precise values come from plasmid sequencing; the success of conjugative transfer is indicated (Transfer of mcr-1). The heat map in the last section indicated the amount of antibiotic resistance genes carried by the respective isolate in specified category of antibiotics from zero (white) to maximum of six (dark purple).

**Figure 2.**
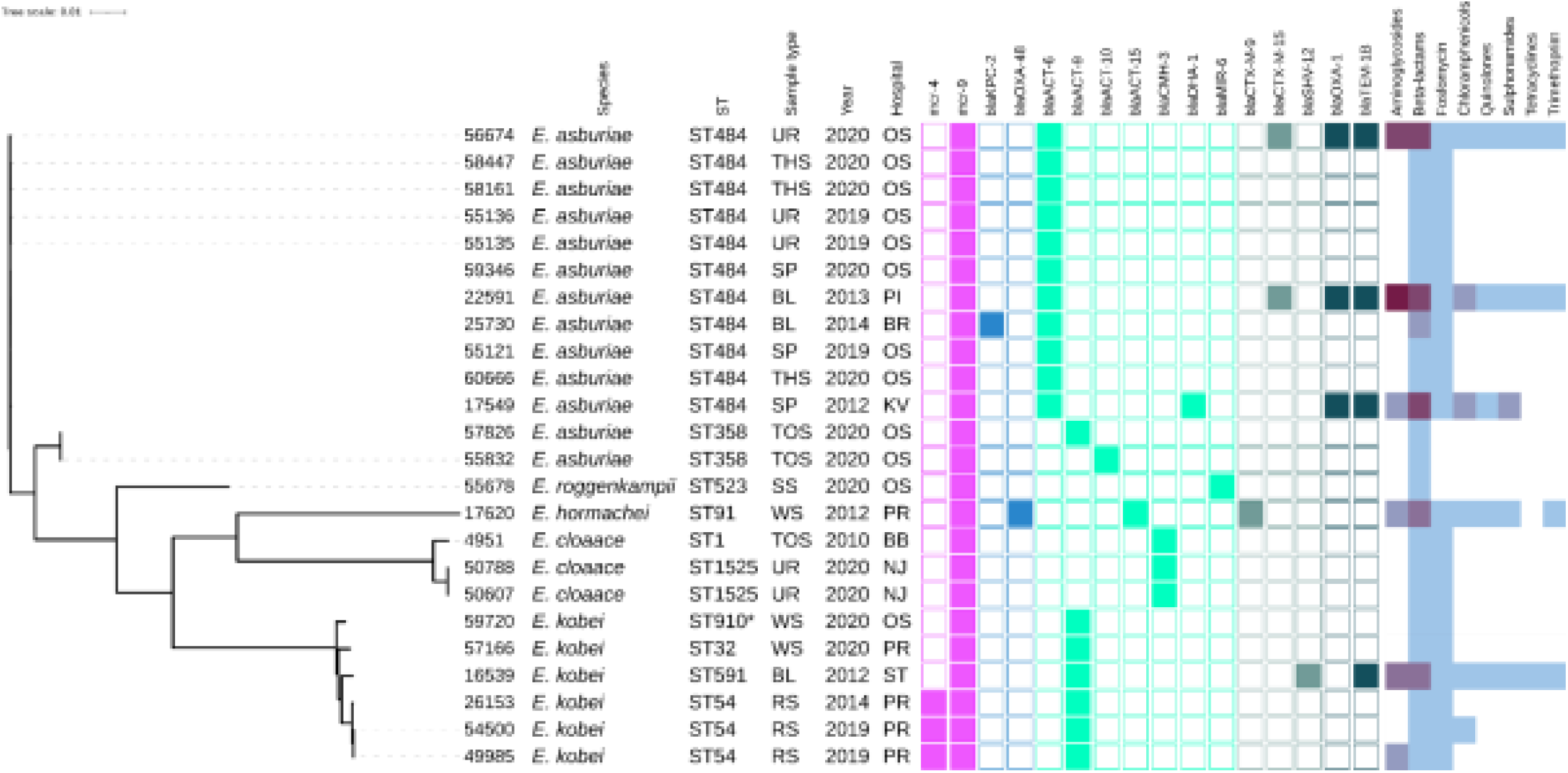
Phylogenetic tree of *Enterobacter* spp. isolates of Czech clinical origin. The metadata specify the species (Species); sequence type (ST), type of sample (Sample type): urine (UR), throat swab (TSH), sputum (SP), blood (BL), tonque swab (TOS), skin swab (SS), wound swab (WS), rectal swab (RS); year of isolation (Year) and city where is the hospital related to the isolate recovery (Hospital): Ostrava (OS), Pribram (PI), Brno (BR), Karlovy Vary (KV), Prague (PR), Brno-Bohunice (BB), Novy Jicin (NJ), Strakonice (ST). The colour squares represent presence (full square) or absence (empty square) or respective antibiotic resistance genes divided to genes encoding resistance to colistin (pink), carbapenemases (blue) and other beta-lactamases (turquoise, see legend Figure 1). The heat map in the last section indicated the amount of antibiotic resistance genes carried by the respective isolate in specified category of antibiotics from zero (white) to maximum of five (dark purple).

Most *mcr-1-*positive isolates carried genes (Supplementary Table S1) conferring resistance to aminoglycosides (36/48), macrolides (42/48), sulphonamides (36/48), tetracycline (33/48) and trimethoprim (31/48). Additionally, 44 *mcr-1*-positive isolates harboured genes for resistance to narrow-spectrum beta-lactams including *bla*_TEM-1B_ (n=27), *bla*_TEM-135_ (n=4) and *bla*_TEM-32_ (n=3). In two isolates, AmpC beta-lactamase genes *bla*_CMY-2_ or *bla*_DHA-1_ were detected while in three isolates, ESBL genes *bla*_CTX-M-1_ (n=1), *bla*_CTX-M-27_ (n=2) were found. All four *mcr-1*-positive *K. pneumoniae* isolates carried *fosA, oqxA, oqxB* and *bla*_SHV_ genes.

On the other hand, the majority of *mcr-9*-positive isolates contained resistance genes to beta-lactams (n=25) and fosfomycin (n=22). Specifically, the *E. asburiae* ST484 isolates carried the *bla*_ACT-6_ gene, encoding the intrinsic AmpC beta-lactamase, while the *E. asburiae* ST358 isolates carried *bla*_ACT-9_ or *bla*_ACT-10_ variants (Figure 2). The two *E. cloacae* ST1525 isolates harboured only *bla*_CMH-3_ gene, encoding the chromosomal AmpC. The ST54 *E. kobei* isolates carried the *bla*_ACT-9_ and *fosA* genes. Moreover, ESBL (*bla*_CTX-M-9_, *bla*_CTX-M-15_) and AmpC beta-lactamase (*bla*_DHA-1_, *bla*_CMY-117_) were identified in three and two *mcr-9*-positive isolates, respectively. One *E. hormaechei* ST91 isolate carried carbapenemase gene *bla*_OXA-48_. The single *Citrobacter freundii* isolate carried five beta-lactamase genes including carbapenemase-encoding gene *bla*_KPC-2_.

Chromosomal mutations to fluoroquinolones (*acrR*), cephalosporins (*ompK36*) and carbapenems (*ompK37*) were identified in four *K. pneumoniae* isolates (Table S1). In contrast with the susceptibility testing, three of these *K. pneumoniae* isolates were resistant to fluoroquinolones and only one to cephalosporins. Twenty-five *E. coli* isolates carried at least one of four mutation variants including *gyrA, gyrB, parC* and *parE* for resistance to quinolones and thirteen *E. coli* isolates carried quinolone-resistance genes *qnrB* (n=5) or *qnrS* (n=8) (Table S1) while MIC profiles showed quinolone resistance in thirty-one isolates. Nine different types of mutations in *pmrA/pmrB* genes associated with colistin resistance were also found in eleven *E. coli* isolates (Table S1).

### Phylogenetic analysis of *E. coli* ST744 and *Enterobacter cloacae* ST484

Phylogenetic analysis of ST744 isolates from our collection (n=10) along with other 38 closely related genomes from public resources showed formation of 2 major clades; four isolates were considered as outliers (Figure 3). First major clade (samples 60462 – 54343, green branch) comprised 17 mostly European isolates from animals and humans, and was further divided into two subclades. Second dominant clade (samples ERR3531597 – ERR1971525, violet branch) was composed of 27 cosmopolitan isolates originating from various sources, and was divided into several smaller subclades. Isolates belonging to different major clades showed a variable number of pairwise SNP differences against each other, ranging from 600 up to 3000. In both major clades, there were apparent clusters of isolates from humans and animals, exhibiting few dozens of SNPs from each other (Supplementary Table S2). Isolates from our collection were scattered across the tree, five of them belonged to the first clade, four to second clade, while one sample was an outlier. Our clinical samples 48907, 52857, 55923 and 54343, belonging to the first clade, showed 46-58 SNP differences from three clinical samples from Germany, and 39-49 SNP differences from the two Romanian (RO) isolates from poultry that carried *bla*_CMY-2_. One of the RO isolates also carried the *mcr-1*.*1* gene that was borne by all isolates from our Czech clinical collection. Those RO isolates were also closely related to three clinical isolates from Germany (exhibited <20 SNP differences from each other). Isolate 60061 from the Czech collection clustered with clinical isolate from Thailand (110 SNPs) and Chinese isolate from a pig (128 SNPs). Notably, the Swiss isolate also carried *mcr-1*.*1*. Our isolates 45082 and 54444 were related to another clinical isolate from the United Kingdom (66 and 73 SNPs) and also to an environmental isolate from a river in Japan (73, and 80 SNPs, respectively). Isolate 52637 from our collection showed the least SNP counts against three Australian isolates from gulls (36-37 SNPs) and three clinical isolates, one coming from Switzerland (36 SNPs), and another from Germany (39 SNPs) and Russia (42 SNPs).

**Figure 3.**
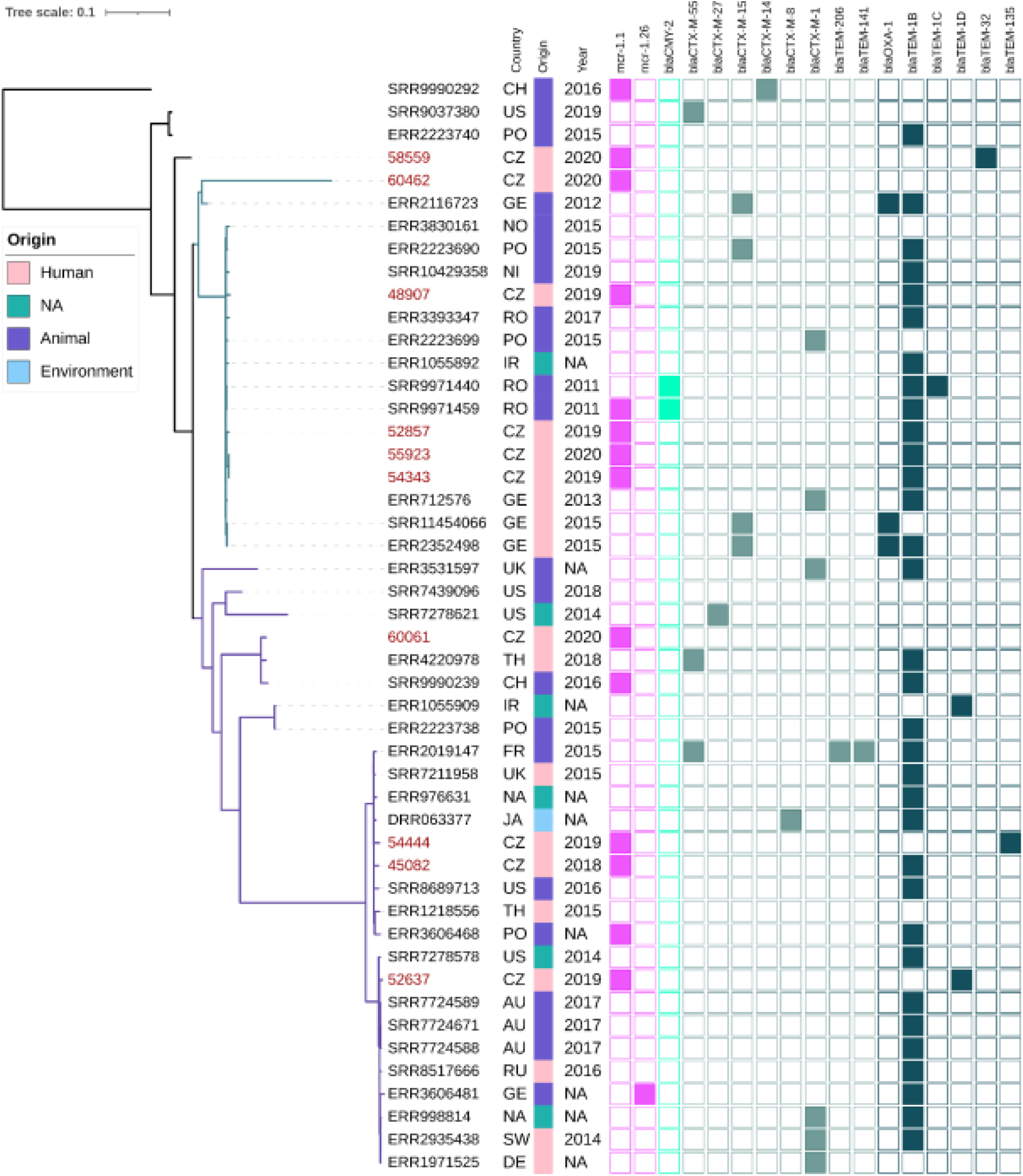
Phylogenetic tree of Czech clinical *E. coli* ST744 isolates with selected sequences from global collection. The red labels indicate isolates coming from this study. The metadata specifies country of origin (Country): China (CH), The United States (US), Poland (PO), Czech Republic (CZ), Germany (GE), Norway (NO), Nigeria (NI), Romania (RO), United Kingdom (UK), Thailand (TH), Ireland (IR), France (FR), Ukraine (UK), Australia (AU), Russia (RU), Switzerland (SW), not available (NA), the source of origin (Source) and the year of isolation (Year). The colour squares represent presence (full square) or absence (empty square) or respective antibiotic resistance genes divided to genes encoding resistance to colistin (pink), carbapenemases (blue) and other beta-lactamases (turquoise, see legend Figure 1).

### Structure of *mcr-1*-carrying plasmids

The *mcr-1* gene was located predominantly on 33 kb IncX4 plasmids (34/48). Six complete plasmids from *E. coli* and *K. pneumoniae* obtained by long-read sequencing (Table 1) showed high level of nucleotide similarity (>99.9%) to each other as well as to plasmids from raw meat from Czech retails (Zelendova et al., 2021) (Supplementary Figure S1). The *mcr-1* gene was bordered by a hypothetical protein and a PAP2 transmembrane protein, which is the typical genetic surrounding for *mcr-1* gene within IncX4 plasmids (Zelendova et al., 2021).

The *mcr-1* was also carried by ∼60 kb IncI2 plasmids (n=8). Plasmid pMCR1-53288 originating from *E. coli* ST538 from urine obtained by MinION sequencing shared high sequence similarity (>98%) with several plasmids available in GenBank database including pMCR_1884_C3 and pMCR_1138_A1 from *C. braakii* and *E. coli* ST162, respectively, isolated from imported meat sold in Czech retails (Zelendova et al., 2021) (Figure S2). The *mcr-1* region was inserted downstream the *nikB* gene, encoding a DNA topoisomerase III, as observed in other IncI2 *mcr-1*-positive plasmids like pMCR_1884_C3. No other resistance genes were located on IncI2 plasmids.

From our collection, six *Enterobacterales* isolates were found to harbour IncHI2 plasmids with *mcr-1* gene. The complete sequence of three *mcr-1* positive IncHI2 plasmids pMCR1-59496, pMCR1-43934 and MCR1-51133 was determined using MinION technology. BlastN analysis showed that all sequenced IncHI2 plasmids, ranging from ∼225 kb to ∼255 kb in size, belonged to ST4 and were closely related (coverage 80-99%, identity 99%) to each other (Figure 4), as well as to other *mcr-1*-carrying IncHI2 plasmids, like pMCR_915_C1 and pMCR_1085_C1 from *E. coli* recovered from imported meat (Zelendova et al., 2021), and plasmid pKP121-1-mcr (Ruan et al., 2019) of human clinical origin from China. All IncHI2 plasmids contained regions for conjugative transfer (*htd, orf, tra* genes) and plasmid maintenance (*par* gene). Additionally, IncHI2 plasmids carried tellurium resistance genes in two clusters including *terZABCDEF* and *terXYW* (except p56099). In all InHI2 plasmids, characterized during this study, *mcr-1* gene was inserted downstream the *terY2* gene, as observed in other IncHI2 plasmids like pMCR_1085_C1. Similar to pMCR_1085_C1, the *mcr-1* gene was bounded by an IS*Apl1* element and PAP2 transmembrane protein (Zelendova et al., 2021). All IncHI2 *mcr-1*-positive plasmids exhibited at least one MDR region, which ranged in size from 950 to 36097 bp (Figure 4).

**Figure 4.**
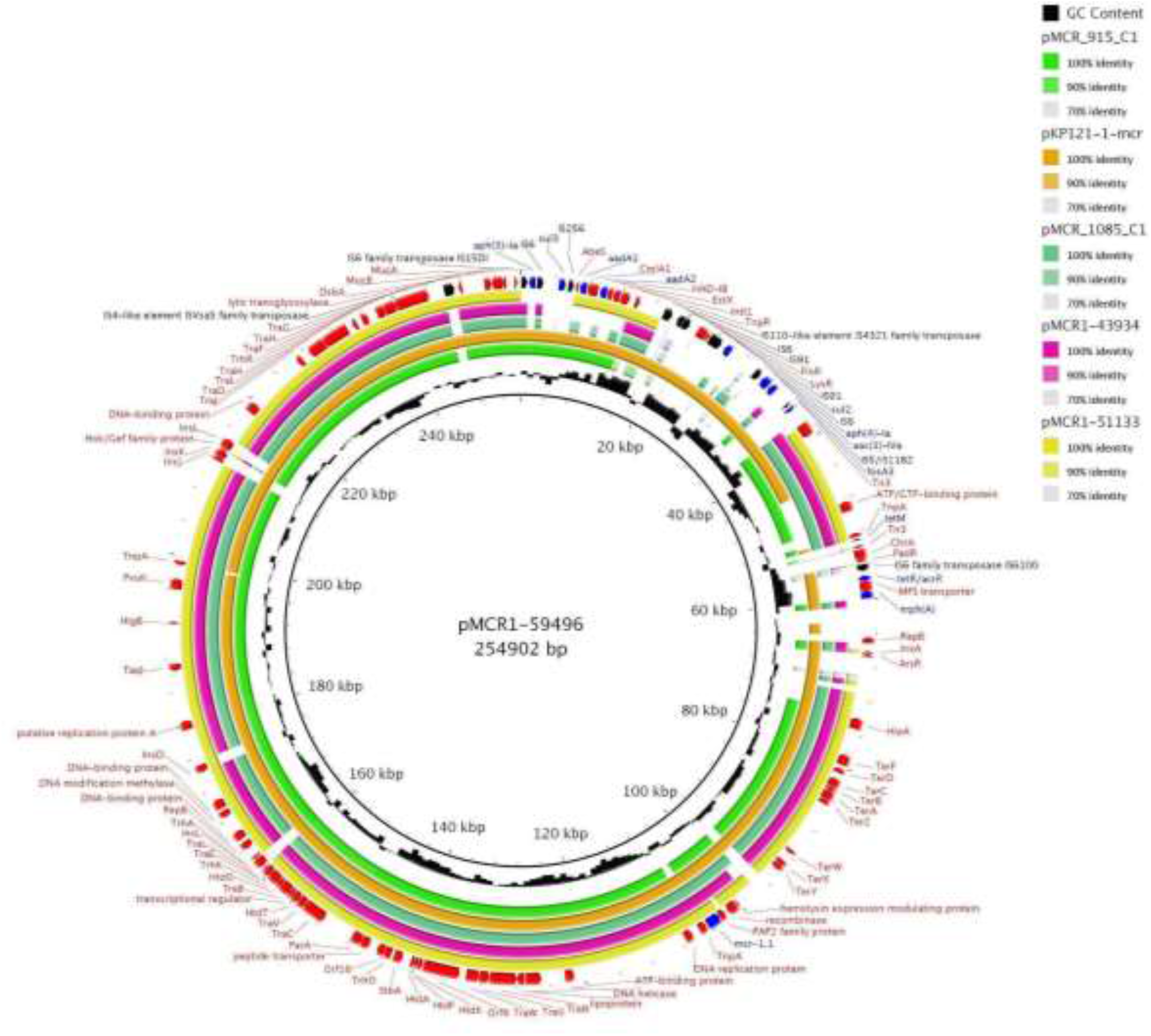
BRIG comparison of *mcr-1*.*1*-encoding IncHI2/ST4 plasmids. The plasmid pMCR1-59496 from *K. pneumoniae* ST726 identified in our study from urine sample (OP950836) was used as a reference. Two other plasmids originated from our collection including pMCR1-43934 from *E. coli* ST8186 from tonsil swab sample and pMCR1-51133 from *E. coli* ST117 from a urine sample. The sequence alignment includes pMCR_915_C1 (MT929284.1) and pMCR_1085_C1 (MT929286.1) from *E. coli* recovered from raw turkey meat imported to the Czech Republic from Poland and one of plasmid pKP121-1-mcr (CP031850.1) from *K. pneumoniae* ST2570 from human blood in China.

### Structure of *mcr-4-*encoding plasmids

*mcr-4* was located on ColE10 plasmids in three *E. kobei* ST54 isolates. Plasmid pMCR4-26153 of size 12,808 kb recovered from a rectal swab of a patient in the Czech Republic, was identical (100% coverage, 100% identity) to pIB2020_ColE_MCR (Marchetti et al. 2021) from *E. kobei* ST54 strain from a rectal swab of a 56 years old male patient hospitalized in 2019 in Italy (Figure S3).

### Structure of *mcr-9*.*1*-carrying elements

Out of the twenty-five *mcr-9*.*1*-positive isolates, eight were characterized by MinION technology. Among the latter isolates, three carried the *mcr-9*.*1* allele on IncHI2 plasmids (Table 1) while, in the five remaining isolates, the *mcr-9*.*1* was found on the chromosome. Plasmid pMCR9-57185 originated from *C. freundii* ST18 recovered from rectal swab while pMCR9-16539 was obtained from *E. kobei* ST591 from blood and pMCR9-17620 came from *E. hormaechei* ST91 recovered from wound swab.

Following the IncHI2 pDLST scheme, plasmids pMCR9-57185 and pMCR9-16539 were typed as ST1, while pMCR9-17620 that differed by a single nucleotide polymorphism in *smr0199* locus was assigned to a novel ST. All plasmids exhibited closely related sequences (>89% coverage, 99.99% identity) to other *mcr-9*.*1*-positive IncHI2 plasmids (Figure S4), like p49790_MCR from an *E. hormaechei* isolate recovered previously from Czech hospitals (Bitar et al., 2020). Similar to p49790_MCR, the *mcr-9*.*1* was inserted upstream the *pcoS* gene (encoding a membrane protein for resistance to copper), in all IncHI2 plasmids like p49790_MCR. Additionally, in plasmids pMCR9-57185 and pMCR9-16539, the *mcr-9*.*1* gene was bounded by an IS element (upstream) and an ORF (downstream), encoding a cupin fold metalloprotein, followed by IS*26*. However, in plasmid pMCR9-17620, an IS*1* was found downstream of *mcr-9*.*1*. Furthermore, IncHI2 plasmids contained at least one MDR region including genes for resistance to aminoglycosides, tetracyclines, trimethoprim, chloramphenicol, sulfonamides, quinolones, and/or macrolides (Table 1). Moreover, IncHI2 plasmids carried tellurium resistance genes (*terZABCDEF*) commonly associated with this plasmid family, and genes conferring arsenic resistance (*arsCBRH*).

The *mcr-9*.*1* gene was integrated into the chromosomes of four *E. cloacae* complex isolates obtained by long-read assembly. The upstream genetic surroundings were identical in all isolates consisting of *mcr-9*.*1*, IS*903B, pcoS, pcoE, rcnA* and *rcnR* genes while the downstream sequences differed. In isolates 50607 and 59720, the *mcr-9*.*1* was followed (downstream) by *wbuC*, IS*26* and IS*1A* forming a region *rcnR*-*rcnA*-*pcoE*-*pcoS*-IS*903B*-*mcr-9*.*1*-*wbuC*-IS*26* identical to the respective region of the plasmid pMCR9-57185. On the other hand, the downstream environment of *mcr-9*.*1* in isolates 56674 and 57166 consisted of *wbuC, qseC, qseB* and ATPase ORF similar to the corresponding region in the chromosome of a Japanese human isolate *Enterobacter asburiae* A2563 (AP022628), visualized in Figure 5.

**Figure 5.**
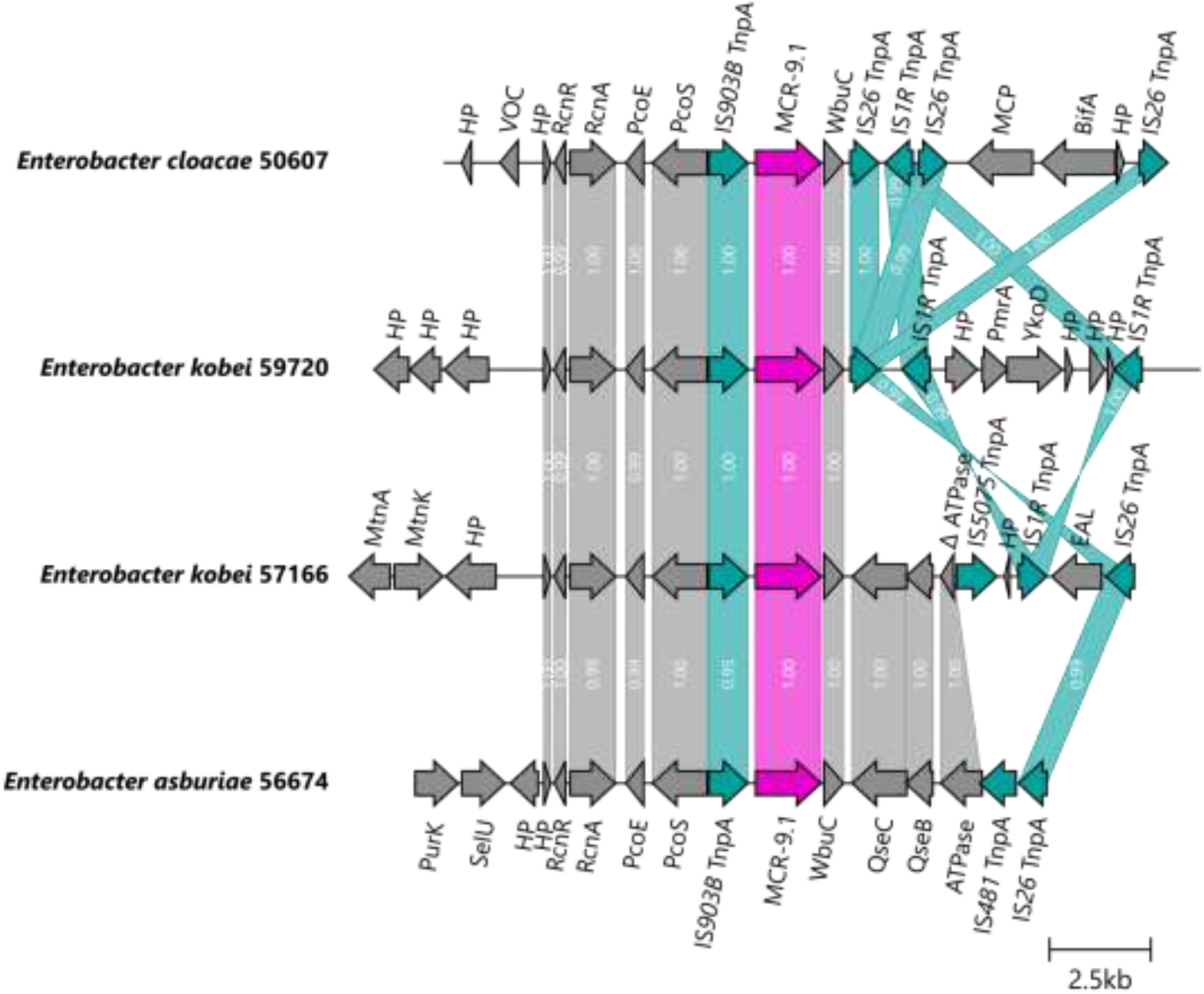
Genetic surroundings of *mcr-9*.*1* gene located on chromosomes in *Enterobacter*. The linearized coding sequences of MCR-9.1 region of four isolates were compared using clinker with identity threshold 90%. The MCR-9.1 (pink) surrounded by mobile genetic elements (turquoise) and other coding sequences (grey) in isolates 50607 and 59720 formed a region corresponding to the *mcr-9* region observed in IncHI2/ST1 plasmids. On the other hand, 57166 and 56674 isolates contained downstream sequence similar to the previously described *mcr-9* environment in *E. asburiae* (AP022628). White numbers in links correspond to the similarity of coding sequences.

### Horizontal transfer of *mcr* gene

Resistance to colistin associated with *mcr-1* was transferred to recipient *E. coli* laboratory strains via conjugation in the majority of *E. coli* isolates (38/44, 86%) and in all isolates of *K. pneumoniae* (4/4, 100%). The most frequently transferred plasmid harbouring *mcr-1* included 33 kb plasmid IncX4 (31/34, 91%), followed by 55 kb IncI2 (8/8, 100%) and 220 kb IncHI2 (4/6, 67%) plasmids. IncHI2 plasmid with *mcr-9* was transferred via conjugation in one *E. asburiae* isolate. Also, ColE10 plasmid carrying *mcr-4* was not transferred via conjugation, since this plasmid family does not contain any transfer region.

### Structure of carbapenemase-encoding plasmids

The *bla*_OXA-48_ was carried by a 63.7 kb IncL plasmid pOXA_17620 (OQ127401) of the *E. hormaechei* ST91 isolate. The plasmid was identical with a previously described plasmid pRIVM_C012525_20 (CP068332). The *bla*_OXA-48_ was surrounded by IS*10A* and IS*1R* upstream and by LysR family transcriptional regulator and IS*10A* downstream (Figure S5).

The *bla*_KPC-2_ was carried by a 50.4 kb IncR plasmid pKPC_57185 (OQ127401) of the *C. freundii* ST18 isolate. The gene was surrounded by Tn*4401* and IS*Kpn7* upstream, and DNA resolvase and Tn*5403* downstream. The pKPC_57185 plasmid was similar (coverage 92%, identity 100%) to a previously described IncR plasmid p46903_KPC (CP070521) (Figure S5). However, the pKPC_57185 carried a MDR region identical with a larger IncN-IncR plasmid (CP070576) from the same study, as visualised in Figure S5.

## Discussion

Within this study, we performed a surveillance of *mcr*-encoding genes among colistin-resistant *Enterobacterales* collected from Czech hospitals between 2009 and 2020. Our findings indicated a low prevalence (3.8%) of *mcr* genes among colistin-resistant isolates with slightly increasing prevalence during the study period (3% in 2018 while 9% in 2020). However, the prevalence of *mcr*-positive isolates may be overestimated since our collection was composed only of colistin-resistant isolates and did not include the bacterial population susceptible to colistin. Moreover, another study limitation is the fact that isolates from retrospective sampling from the period 2009-2017 were obtained during various surveillance programs at the National Institute of Public Health. As these programs were focused mainly on *Klebsiella* sp, invasive *E. coli* or they were targeting MDR strains, the data of *mcr* prevalence from this period needs to be interpreted with caution.

Another Czech study (Tkadlec et al., 2021) published a low prevalence of the *mcr-1* gene (4/1922, 0.21%) in *E. coli* from fecal samples from hospitalized patients between June 2018 and September 2019. A study from Switzerland reported that the fecal carriage rate of colistin-resistant (MIC value >2 mg/l) *Enterobacterales* was 1.5% for healthy people and 3.8% for primary care patients, while none of the isolates harboured the *mcr-1* or *mcr-2* genes (Zurfluh et al. 2017). Additionally, in Finland, only one *mcr-1*-positive *E. coli* was characterized from fecal samples collected from 176 healthy volunteers (Kirsi Gröndahl-Yli-Hannuksela 2018), during 2016. Other studies from Europe have reported low prevalence of colistin-resistant isolates and of *mcr*-positive *Enterobacterales*. In Spain, the overall prevalence of colistin resistance in clinical isolates of *Enterobacterales* was 0.67%. The rate was higher in *E. cloacae* (4.2%) than *E. coli* (0.5%) and *K. pneumoniae* (0.4%) while *mcr-1* was detected only in *E. coli* (0.15%) (Prim et al. 2017). Similar prevalence levels were observed for Romagna, Northern Italy, where the prevalence of colistin-resistant isolates among human *Enterobacterales* was 0.5% and the *mcr-1* gene was found in 0.14% *E. coli* isolates (Bianco 2018). On the other hand, higher percentages have been reported regarding the prevalence of colistin-resistant isolates in different geographical areas. Giani et al. (2018) reported a high proportion (38.8%) of *mcr-1* carriers among healthy children (129/337) from Bolivia. Furthermore, in Chinese hospitals across 24 provincial capital cities and municipalities, human carriage of *mcr-1*-positive *E coli* was identified in 644 (14.3%) of 4498 samples in 2016 (Wang 2020). However, different methodological approaches and study design (e.g., selective cultivation on colistin-supplemented media, targeting isolates despite their susceptibility profiles, PCR detection of *mcr* genes in either total enterobacterial microbiota or directly in a clinical sample) significantly limit the comparison of prevalence data between the studies. Moreover, the discrepancy in the prevalence of *mcr* carriers between studies and geographical regions underlines the other factors, like antibiotic use and stewardship protocols, contributing in the emergence and spread of colistin-resistant isolates.

In this study, we found *pmrA* or *pmrB* genes mutations associated with chromosomal colistin resistance in *E. coli* isolates carrying *mcr* (11/44, 25%). However, our collection also contained twenty-four *Enterobacter* spp. isolates and one *C. freundii* in which colistin resistance is often associated with mutations in two-component regulator systems PmrAB and PhoPQ (Hong and Ko, 2019; Wand and Sutton, 2020) and these mutations were not the target of the study. Majority of those isolates carried *mcr-1* allele (n=48) while twenty-two isolates harboured the *mcr-9*.*1* allele and the three remaining isolates co-carried the *mcr-4*.*2*/*mcr-4*.*3* and *mcr-9*.*1* genes. These results are consistent with the current global epidemiology of *mcr* genes where *mcr-1* and *mcr-9* are most widely disseminated (Ling et al., 2020). Most *mcr-1* carriers were *E. coli* and as the gene was present in 23% of all resistant isolates of this species, which is in agreement with findings of previous study (Zelendova et al., 2021). Additionally, a retrospective screen of colistin-resistant *Enterobacterales* reported in the National Institute of Public Health from Czech hospitals, during 2009-2017, revealed the presence of nine additional isolates carrying *mcr* genes. Eight of the latter isolates produced MCR-9, whereas the *E. kobei* ST54 strain also expressed the MCR-4 protein. The remaining isolate, a *K. pneumoniae* ST2590 isolated in 2017, produced MCR-1.1 protein. Most of *mcr-9*.*1*-carrying isolates belonged to *Enterobacter* spp. Previous studies have shown that *mcr-9* gene is commonly associated with isolates belonging to *Salmonella* and *Enterobacter* genus (Zhang 2022; Liao 2022; Bitar et al., 2020). Interestingly, the low resistance levels to colistin of MCR-9-producing *Enterobacter* isolates has been reported (Bitar et al., 2020). This observation may explain the unnoticed spread of those isolates in Czech hospitals. Remarkably, 96% of isolates (70/73) carried AmpC/ESBL or carbapenemases, raising the concern that the spread of *mcr-*carrying isolates might also be related to the use of other antimicrobial agents including clinically important beta-lactams. IncHI2 plasmids carrying *mcr-9*.*1* harboured also genes for resistance to aminoglycosides, beta-lactams, trimethoprim, sulphonamides and/or tetracyclines (Table 1).

MLST revealed the presence of *mcr* genes in various STs of *E. coli, K. pneumoniae*, and *Enterobacter* sp., highlighting the significant impact of horizontal gene transfer in the spread of colistin resistance determinants via plasmids. Phylogenetic analysis of *E. coli* ST744 isolates, the dominant *E. coli* genotype, showed formation of 2 major clades (Figure 3). Five isolates from our collection belonged to the first clade, which comprised mostly European isolates from animals and humans, were closely related with three clinical isolates from Germany and two samples of poultry origin from Romania. The second dominant clade contained four isolates from our collection, which were closely related with isolates from different geographical areas and various sources (Figure 3). Additionally, phylogenetic analysis uncovered the association of *mcr* genes with specific clones, like *E. kobei* ST54, which has been previously reported to produce MCR-4.3 from clinical samples recovered in Italy (Marchetti et al., 2021). Of note, these observations underline the important role of travelling across the borders, that has contributed to the spread of MDR bacteria.

Finally, analysis of *mcr*-carrying plasmid sequences showed the presence of *mcr-1*, mainly on IncX4 replicons, but also on IncI2 and IncHI2 plasmids. On the other hand, the *mcr-9* allele was found on IncHI2 plasmids (n=3) or it was integrated into the chromosome of *Enterobacter* isolates (n=5). The *mcr-4* gene was located on ColE10 plasmids. These findings are in agreement with the previously published data, showing the emergence of *mcr* genes on the specific Inc groups of plasmids that were characterized from *Enterobacterales* recovered from different sources including animals, food and humans (El Garch et al., 2018; Xavier et al., 2016; Zurfluh et al., 2017; Bitar et al., 2020; Li et al., 2020; Zelendova et al., 2021; Marchetti et al., 2021). Furthermore, our experiments demonstrated a high efficiency of conjugative transfer of *mcr-1*-carrying IncX4 plasmids. Also, the conjugative transfer of IncHI2 plasmids carrying *mcr-1* or *mcr-9* was confirmed. Thus, the horizontal transfer of plasmid-mediated *mcr* genes represents an important risk factor for public health since colistin is considered as one of the last-resort antibiotics for the treatment of serious infections in human medicine. Therefore, studying the spread of MDR pathogens is vital for analysis of transmission pathways and risk factors for public health.

The prospective epidemiological survey performed in this study brought the first information on the plasmid-mediated dissemination in the Czech Republic and showed that a surveillance system is essential to monitor the diffusion of plasmid mediated colistin resistance.

---

**Figure S1.**
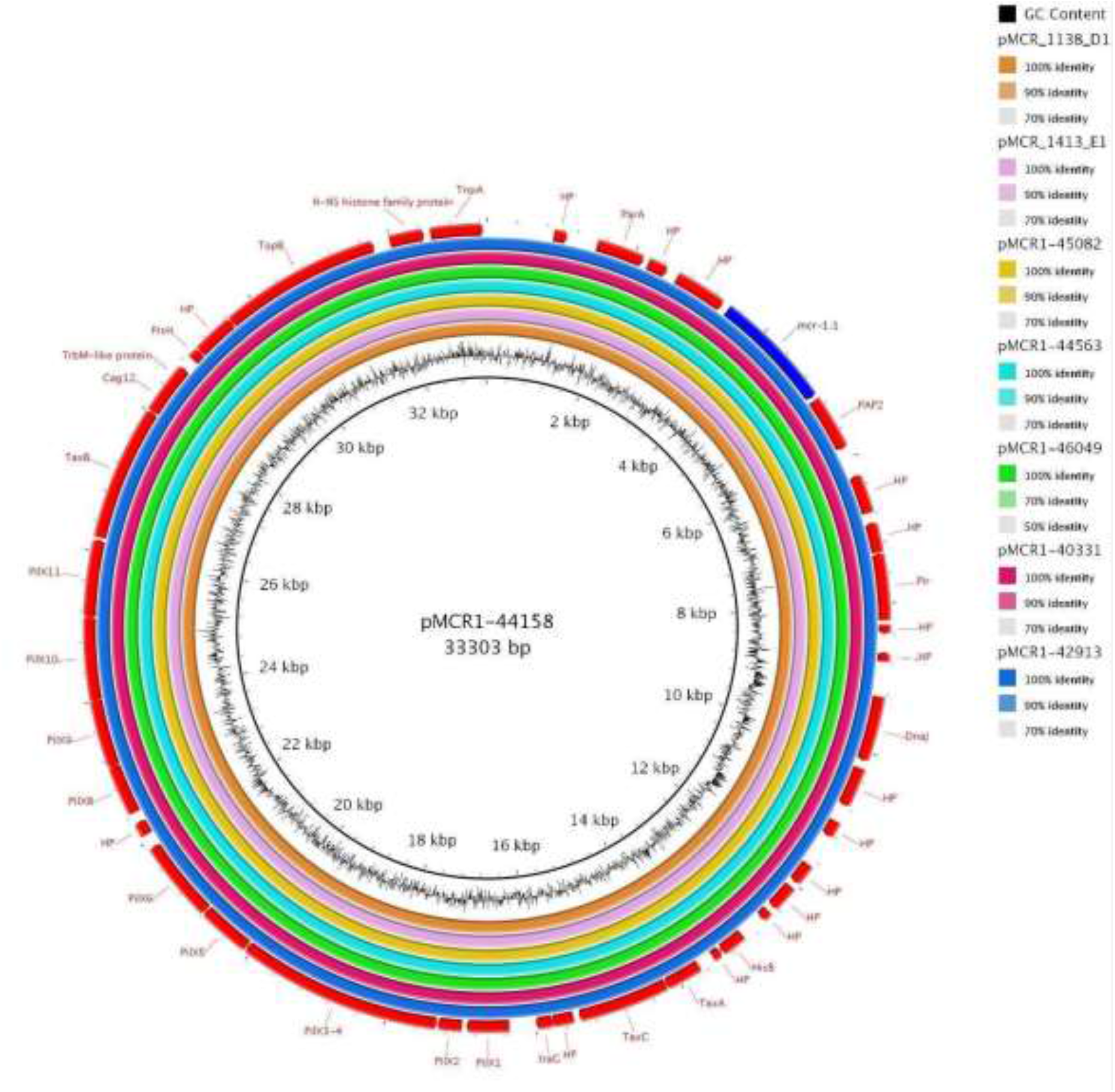
BRIG comparison of *mcr-1-*encoding IncX4 plasmids. Six representative plasmids from our study subjected to MinION sequencing were used in the alignment. The plasmid pMCR1-44158 carrying *mcr-1*.*2* from *E. coli* ST156 recovered from wound swab (OP428975) was used as a reference for the comparison. Other plasmids originated from *E. coli* of different STs including pMCR1-42913 (from *E. coli* ST448, rectal swab), pMCR1-44563 (from *E. coli* ST1196, urine) and pMCR1-45082 (from *E. coli* ST744, blood). Other two plasmids from our study came from *K. pneumoniae* including pMCR1-40331 (*K. pneumoniae* ST290, urine) and pMCR1-46049 (*K. pneumoniae* ST147, pus). The sequence alignment contains plasmid sequences from other sources including pMCR_1413_E1 (MT929275) from *E. coli* ST354 from Czech raw turkey meat and pMCR_1138_D1 (MT929276) from *E. coli* ST744 from raw turkey meat imported to the Czech Republic from Germany.

**Figure S2.**
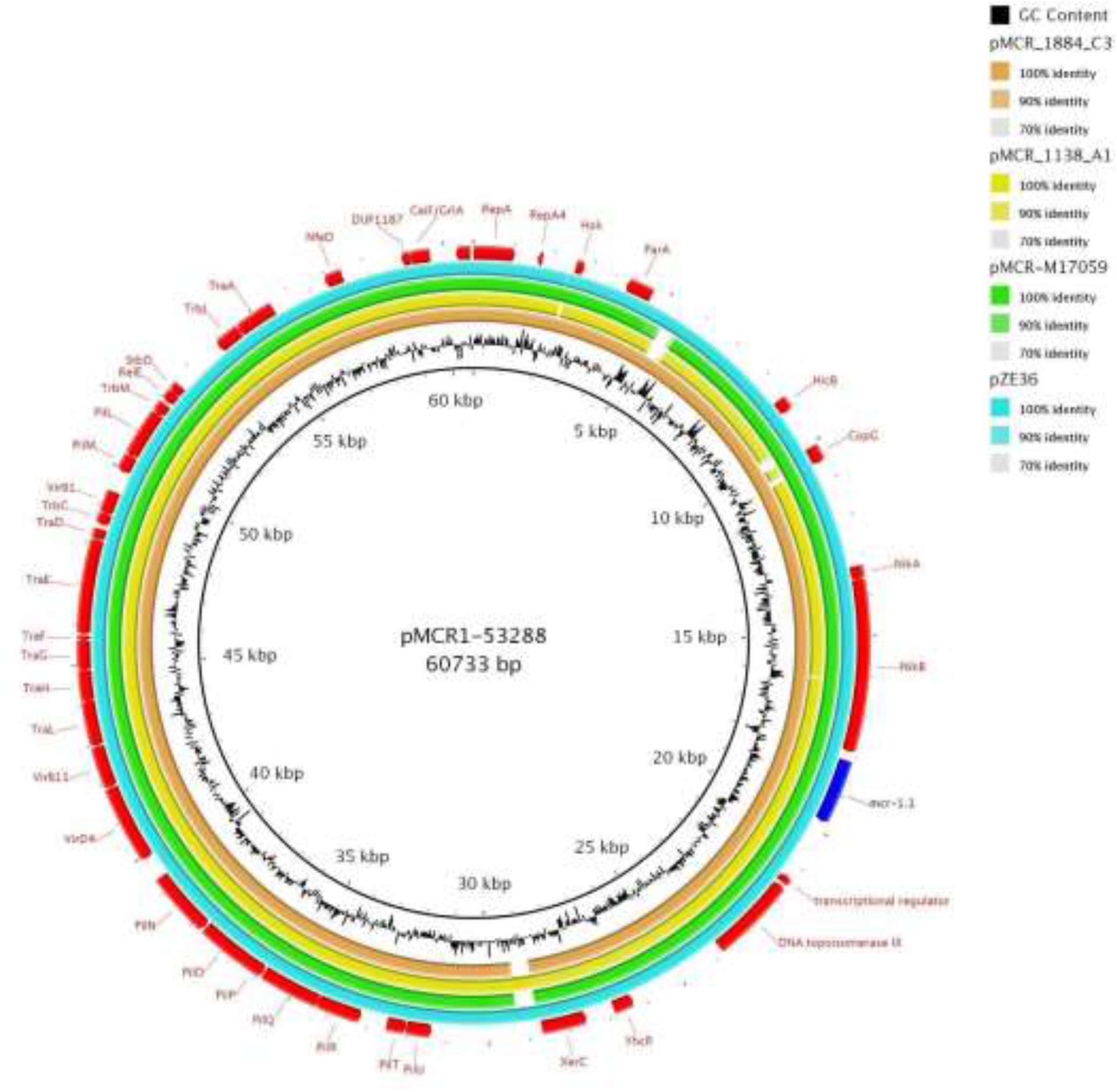
BRIG comparison of *mcr-1*.*1*-encoding IncI2 plasmids. Plasmid pMCR1-53288 originating from *E. coli* ST538 recovered from a urine sample (OP434482) from our study was used as a reference. Sequence alignment contains pMCR_1884_C3 (MT929290) identified in *C. braakii* from raw rabbit meat imported to the Czech Republic from China, while pMCR_1138_A1 (MT929289) originated from *E. coli* ST162 from raw turkey meat imported to the Czech Republic from Germany. The plasmids pMCR-M17059 (KY471310) and pZE36 (KY802014), both of clinical origin, were obtained from *E. coli* ST1488 from Argentina and from *E. coli* ST156 from China.

**Figure S3.**
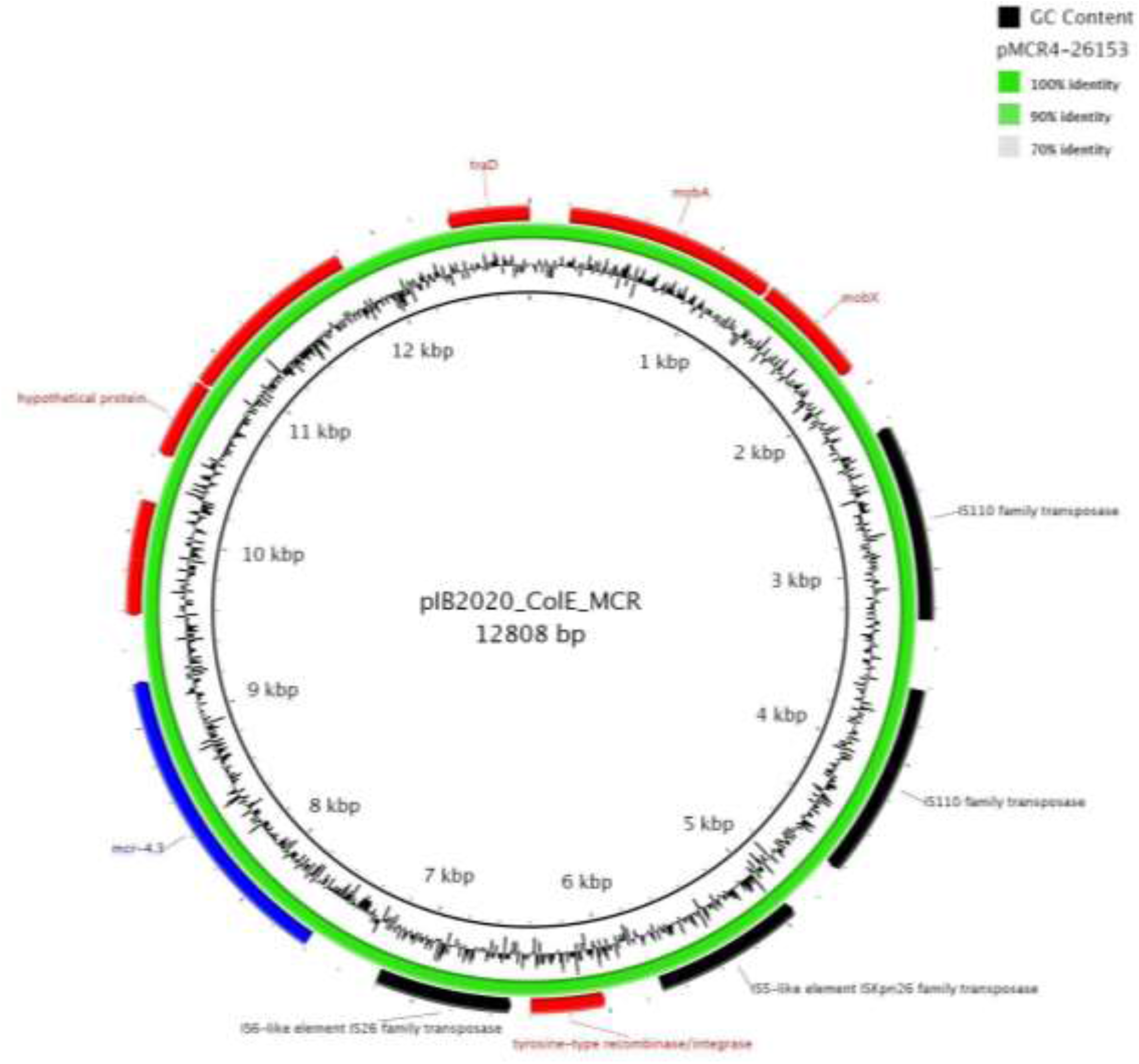
BRIG comparison of ColE10 plasmids with *mcr-4*. pMCR4-26153 originates from *E. kobei* ST54 recovered from a rectal swab of a patient in the Czech Republic. Plasmid pIB2020_ColE_MCR (CP059482) from *E. kobei* ST54 from a patient in Italy was used as a reference.

**Figure S4.**
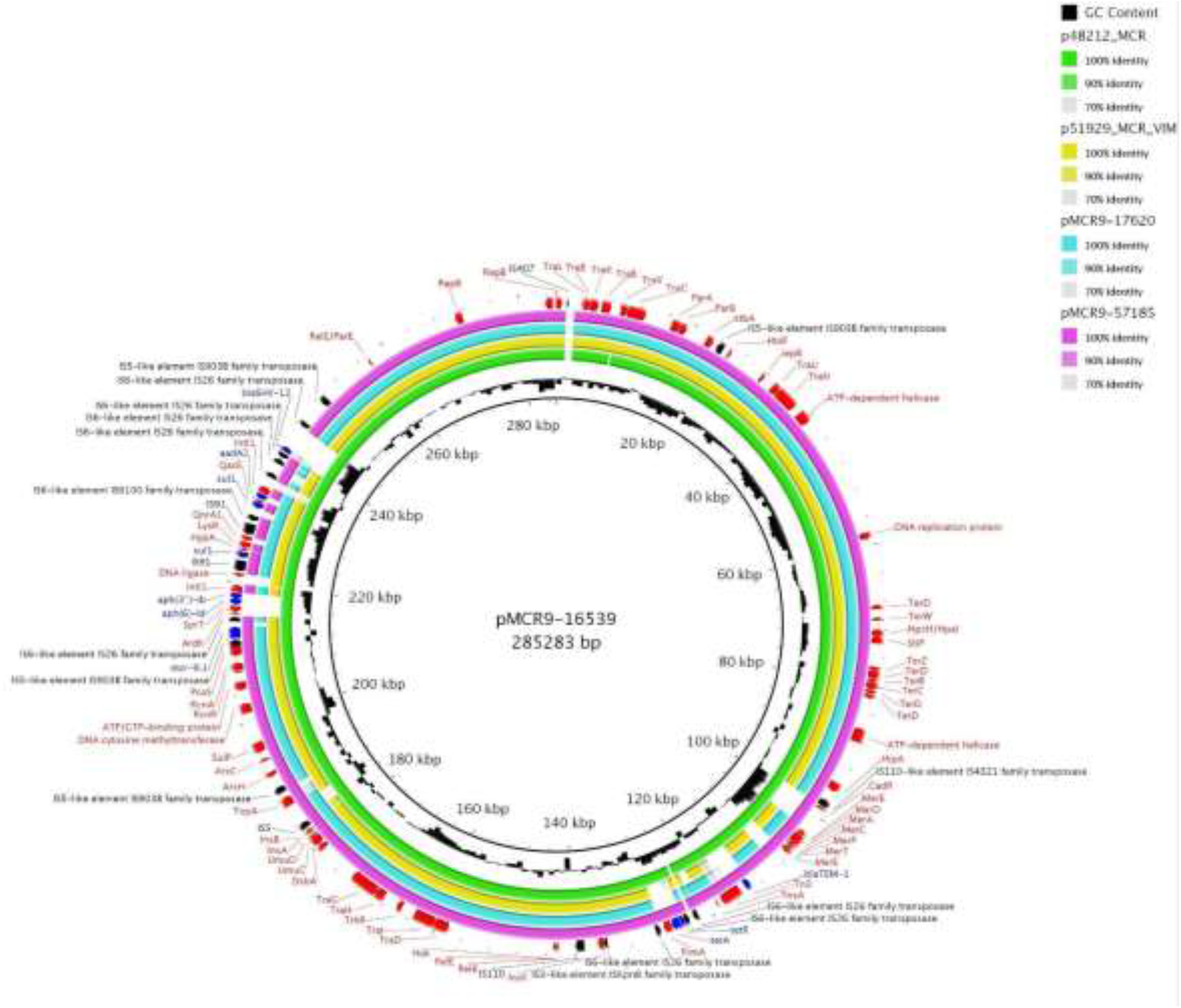
BRIG comparison of *mcr-9*.*1-*encoding IncHI2/ST1 plasmids. The plasmid pMCR9-16539 from *E. kobei* ST591 recovered from a blood sample (OP950838) from our collection was used as a reference. Other two plasmids from our study included pMCR9-57185 (*C. freundii* ST18, rectal swab) and pMCR9-17620 (*E. hormaechei* ST91, wound swab). Plasmids from human clinical isolates previously characterized from the Czech Republic including *C. freundii* (p51929_MCR_VIM; CP059429) and *E. hormaechei* (p48212_MCR; CP059413) were used for the comparison.

**Figure S5.**
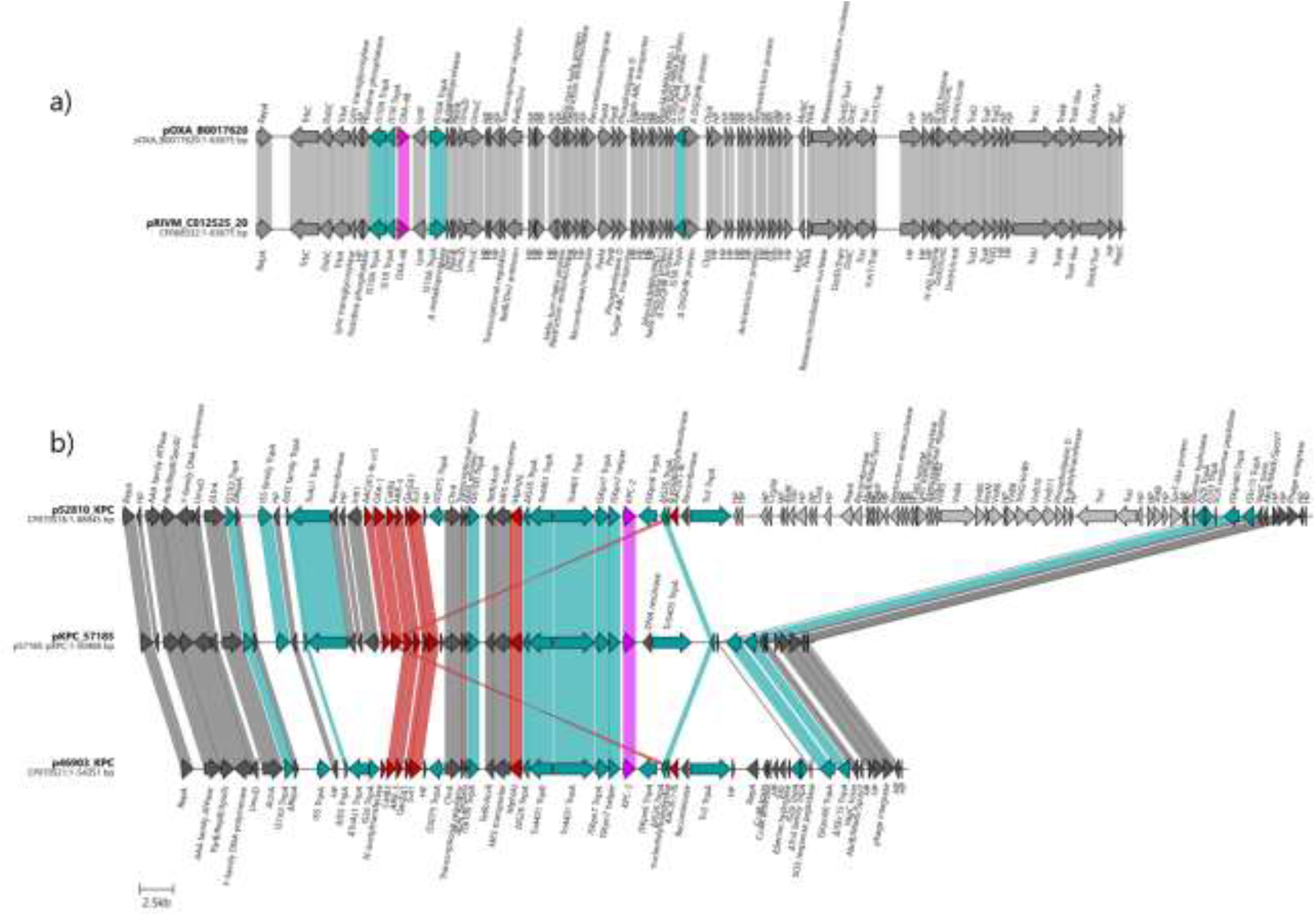
Genetic comparison of the carbapenemase-encoding plasmids. The linearized coding sequences of the carbapenemase-encoding plasmids were compared with reference plasmids using clinker with identity threshold 90%. The coding sequences of OXA-48 and KPC-2 are visualised in pink and other antibiotic resistance CDS in dark red. Turquoise shading indicates mobile genetic elements (MGEs) and grey suggests other coding sequences. HP stands for hypothetical protein. a) The coding sequences of the pOXA_17620 were identical with a reference plasmid pRIVM_C012525_20 (CP068332). b) The linearized coding sequences of the pKPC_57185 were compared with reference plasmids p52810_KPC (CP070576) and p46903_KPC (CP070521). The pKPC_57185 carried multiple MGEs and antimicrobial resistance genes, similarly as the reference plasmid p52810_KPC. The downstream sequence of the KPC-2 differs in the studied plasmid. The KPC-2 is followed by DNA resolvase and Tn*5403*, unlike the reference plasmids carrying IS*Kpn6*, part of IS*26*, AAC(6’)-Ib and recombinase followed by Tn*3*. Other IncR and IncN coding sequences are visualised in dark grey and light grey, respectively.

## Author Contributions

MZ performed laboratory work, data analysis and prepared the manuscript. CCP performed data analysis and prepared the manuscript. PS performed laboratory work and helped with the manuscript preparation. MM performed bioinformatic analyses of whole-genome sequencing data. JP conducted MinION sequencing and helped with plasmid analysis and KN contributed on figure preparation. VJ, KP and HZ provided the samples and IJ performed whole-genome sequencing. MD supervised the project, performed data analysis and revised the manuscript. All authors discussed the results. Members of the surveillance network provided the clinical isolates and metadata obtained during the standard microbiological testing in their laboratories.

## Working Group for Monitoring of Antibiotic Resistance

Vaclava Adamkova, First Faculty of Medicine and University Hospital, Charles University, Prague; Natasa Bartonikova,Bata’s Hospital, Tamara Bergerova, Faculty of Medicine and University Hospital in Plzen, Charles University, Plzen; Marie Bohackova, Hospital in Chrudim, Chrudim; Czysova Erika, Hospital in Sumperk, Sumperk; Josef Cermak, Health institute Usti nad Labem, Kladno; Martina Curdova, Military Hospital Praha, Daniela Fackova, Liberec Regional Hospital, Liberec; Linda Drabkova, University Hospital in Brno, Brno; Lenka Dvorakova, Masaryk Hospital in Usti nad Labem, Usti nad Labem; Galina Eliasova, Regional Hospital Kladno, Kladno; Vladimir Fibiger, Hospital and polyclinic Ceska Lipa, Ceska Lipa; Marian Glasnak, Rudolf and Stefania Benesov Hospital, Benesov; Vera Haskova, Health institute Usti nad Labem, Horovice; Gabriela Hedvicakova, Hospital in Semily, Semily; Blanka Horova, Bulovka University Hospital, Prague; Eva Chmelarova, AGELLAB, Ostrava-Vitkovice; Jan Kubele, Hospital Na Homolce, Prague; Eva Jechova, Thomayer University Hospital, Prague; Petr Jezek, Regional Hospital in Pribram, Pribram; Helena Jordakova, University Hospital in Kralovske Vinohrady, Prague; Jana Jurankova, SPADIA LAB, Brno; Miloslava Kocianova, SYNLAB, Prague; Ivana Kohnova, AGEL Prostejov Hospital, Prostejov; Dana Krckova, IFCOR-99, Brno; Eva Krejci, Health institute in Ostrava, Ostrava; Hana Kremeckova, Hospital in Kyjov, Kyjov; Alice Kucharova, Hospital in Tabor; Katerina Laskafeldova, AGEL Laboratory, Novy Jicin; Jiri Malina, AeskuLab Hadovka, Prague; Eva Martinkova, DIA-GON MP, Cheb; Monika Mazurova, Hospital Usti nad Labem, Zamberk; Marian Mednansky, Hospital in Havlickuv Bod, Havlickuv Brod; Eliska Miskova, Hospital in Trebic, Trebic; Lenka Nanakova, Hospital in Hodonin, Hodonin; Helena Nedvedova, Hospital in Klatovy, Klatovy; Otakar Nyc, University Hospital in Motol, Charles University, Prague; Blanka Ochvatova, SPADIA LAB, Ostrava; Pavla Paterova, University Hospital in Hradec Kralove, Hradec Kralove; Zdena Pitakova, Hospital in Vyskov; J. Podrouzkova, Sang Lab, Karlovy Vary; Miroslava Prejzkova, Synlab, Chomutov; Renata Pribikova, Hospital in Litomerice; Blanka Puchalkova, Hospital in Karlovy Vary, Karlovy Vary; Jana Repiscakova, Hospital in Uherske Hradiste, Uherske Hradiste; Zuzana Semerakova, SPADIA LAB, Prague; Helena Skacani, Hospital in Jihlava, Jihlava; Marketa Skruzna, Institute of Clinical and Experimental Medicine, Prague; Marie Smolikova, Hospital in Jicin, Jicin; Martina Sosikova, Silesian Hospital in Opava, Opava; Michal Stanek, Hospital in Znojmo, Znojmo; Alena Steinerova, CITYLAB, Prague; Petra Safarova, Laboratory of Medical Microbiology, Pardubice; Lenka Semberova, Czech laboratory, Prague; Eva Simeckova, Hospital in Strakonice, Strakonice; Ljuba Suchmanova, Health institute Usti nad Labem, Plzen; David Sus, Hospital in Ceske Budejovice, Ceske Budejovice; Renata Tejkalova, University Hospital of St. Anna, Masaryk University, Brno; Jan Tkadlec, Hospital in Vsetin, Vsetin; Lenka Unuckova, Hospital in Kolin, Kolin; Danuta Urbusova, AGEL Laboratory, Trinec; Vera Kurkova, Hospital in Pisek, Pisek; Denisa Vesela, Hospital in Jindrichuv Hradec, Jindrichuv Hradec; Eva Vesela, Hospital in Nachod, Nachod; Eva Vitova, Hospital in Trutnov, Trutnov; Eva Zalabska, Hospital in Pardubice, Pardubice; Dana Zamazalova, Hospital in Nove Mesto Na Morave, Nove Mesto Na Morave; Roman Zaruba, Hospital in Most, Most; Ilona Zemanova, VIDIA DIAGNOSTIKA, Prague

## Acknowledgments

We thank Dana Cervinkova, Iva Sukkar and Katarina Stredanska for their assistance in the laboratory and to Adam Valcek for his help with data analysis.

## References

Aghapour, Z., Gholizadeh, P., Ganbarov, K., Bialvaei, A. Z., Mahmood, S.S., Tanomand, A., Yousefi, M., Asgharzadeh, M., Yousefi, B. and Kafil, H.S. (2019). Molecular mechanisms related to colistin resistance in Enterobacteriaceae. Infection and Drug Resistance, 12, 965–975. doi:10.2147/IDR.S199844.

Alikhan, N. F., Petty, N. K., Ben Zakour, N. L. and Beatson, S. A. (2011). BLAST Ring Image Generator (BRIG): simple prokaryote genome comparisons. BMC Genomics, 12:1–10. doi:10.1186/1471-2164-12-402.

Bankevich, A., Nurk, S., Antipov, D., Gurevich, A.A., Dvorkin, M., Kulikov, A.S., Lesin, V.M., Nikolenko, S.I., Pham, S., Prjibelski, A.D., Pyshkin, A.V., Sirotkin, A.V., Vyahhi, N., Tesler, G., Alekseyev, M.A. and Pevzner, P.A. (2012). SPAdes: A New Genome Assembly Algorithm and Its Applications to Single-Cell Sequencing. Journal of Computational Biology, 19(5), 455–477. doi:10.1089/cmb.2012.0021.

Bauer, A.P., Dieckmann, S.M., Ludwig, W. and Schleifer, K.-H. (2007). Rapid identification of Escherichia coli safety and laboratory strain lineages based on Multiplex-PCR. FEMS Microbiology Letters, 269(1), 36–40. doi:10.1111/j.1574-6968.2006.00594.x.

Bitar, I., Papagiannitsis, C.C., Kraftova, L., Chudejova, K., Mattioni Marchetti, V. and Hrabak, J. (2020). Detection of Five mcr-9 -Carrying Enterobacterales Isolates in Four Czech Hospitals. mSphere, 5(6). doi:10.1128/msphere.01008-20.

Bolger, A.M., Lohse, M. and Usadel, B. (2014). Trimmomatic: a flexible trimmer for Illumina sequence data. Bioinformatics, 30(15), 2114–2120. doi:10.1093/bioinformatics/btu170.

Borowiak, M., Hammerl, J.A., Deneke, C., Fischer, J., Szabo, I. and Malorny, B. (2019). Characterization of mcr-5-Harboring Salmonella enterica subsp. enterica Serovar Typhimurium Isolates from Animal and Food Origin in Germany. Antimicrobial Agents and Chemotherapy, 63(6). doi:10.1128/aac.00063-19.

Cai, J., Cheng, Q., Shen, Y., Gu, D., Fang, Y., Chan, E.W.-C. and Chen, S. (2017). Genetic and Functional Characterization of blaCTX-M-199, a Novel Tazobactam and Sulbactam Resistance-Encoding Gene Located in a Conjugative mcr-1-Bearing IncI2 Plasmid. Antimicrobial Agents and Chemotherapy, 61(7). doi:10.1128/aac.00562-17.

Camacho, C., Coulouris, G., Avagyan, V., Ma, N., Papadopoulos, J., Bealer, K. and Madden, T.L. (2009). BLAST+: architecture and applications. BMC Bioinformatics, 10(1), 1–9. doi:10.1186/1471-2105-10-421.

Carattoli, A., Bertini, A., Villa, L., Falbo, V., Hopkins, K.L. and Threlfall, E.J. (2005). Identification of plasmids by PCR-based replicon typing. Journal of Microbiological Methods, 63(3), 219–228. doi:10.1016/j.mimet.2005.03.018.

Carattoli, A., Zankari, E., García-Fernández, A., Voldby Larsen, M., Lund, O., Villa, L., Møller Aarestrup, F. and Hasman, H. (2014). In SilicoDetection and Typing of Plasmids using PlasmidFinder and Plasmid Multilocus Sequence Typing. Antimicrobial Agents and Chemotherapy, 58(7), 3895–3903. doi:10.1128/aac.02412-14.

Carattoli, A., Villa, L., Feudi, C., Curcio, L., Orsini, S., Luppi, A., Pezzotti, G. and Magistrali, C.F. (2017). Novel plasmid-mediated colistin resistance mcr-4 gene in Salmonella and Escherichia coli, Italy 2013, Spain and Belgium, 2015 to 2016. Eurosurveillance, 22(31). doi:10.2807/1560-7917.es.2017.22.31.30589.

Caspar, Y., Maillet, M., Pavese, P., Francony, G., Brion, J.-P., Mallaret, M.-R., Bonnet, R., Robin, F., Beyrouthy, R. and Maurin, M. (2017). mcr-1 Colistin Resistance in ESBL-Producing Klebsiella pneumoniae, France. Emerging Infectious Diseases, 23(5), 874–876. doi:10.3201/eid2305.161942.

Centers for Disease Control and Prevention (CDC) (2004). Standardized molecular subtyping of foodborne bacterial pathogens by pulse-field gel electrophoresis. Centers for Disease Control and Prevention, Atlanta, GA.

Dalmolin, V. T., de Lima-Morales, D., Barth, L.A. (2018). Plasmid-mediated Colistin Resistance: What Do We Know? Journal of Infectiology, 1(2), 16–22. doi:10.29245/2689-9981/2018/2.1109.

Darling, A.E., Mau, B. and Perna, N.T. (2010). progressiveMauve: Multiple Genome Alignment with Gene Gain, Loss and Rearrangement. PLoS ONE, 5(6), e11147. doi:10.1371/journal.pone.0011147.

D’Onofrio, V., Conzemius, R., Varda-Brkić, D., Bogdan, M., Grisold, A., Gyssens, I.C., Bedenić, B. and Barišić, I. (2020). Epidemiology of colistin-resistant, carbapenemase-producing Enterobacteriaceae and Acinetobacter baumannii in Croatia. Infection, Genetics and Evolution, 81, 104263. doi:10.1016/j.meegid.2020.104263.

Doumith, M., Godbole, G., Ashton, P., Larkin, L., Dallman, T., Day, M. and Johnson, A. P. (2016). Detection of the plasmid-mediated mcr-1 gene conferring colistin resistance in human and food isolates of Salmonella enterica and Escherichia coli in England and Wales. Journal of Antimicrobial Chemotherapy, 71(8), 2300–2305. doi:10.1093/jac/dkw093

ECDC Technical report. (2019). ISBN: 978-92-9498-300-8.

El Garch, F., de Jong, A., Bertrand, X., Hocquet, D. and Sauget, M. (2018). mcr-1-like detection in commensal Escherichia coli and Salmonella spp. from food-producing animals at slaughter in Europe. Veterinary Microbiology, 213, 42–46. doi:10.1016/j.vetmic.2017.11.014.

El-Sayed Ahmed, M.A.E.-G., Zhong, L.-L., Shen, C., Yang, Y., Doi, Y. and Tian, G.-B. (2020). Colistin and its role in the Era of antibiotic resistance: an extended review (2000–2019). Emerging Microbes & Infections, 9(1), pp.868–885. doi:10.1080/22221751.2020.1754133.

The European Committee on Antimicrobial Susceptibility Testing. 2017. Breakpoint tabled for interpretation of MICs and zone diameters. Version 2.0. http://www.eucast.org.

European Committee for Antimicrobial Susceptibility Testing (EUCAST) of the European Society of Clinical Microbiology and Infectious Diseases (ESCMID). Determination of minimum inhibitory concentrations (MICs) of antibacterial agents by broth dilution. (2003). Clinical Microbiology and Infection, 9(8), doi:10.1046/j.1469-0691.2003.00790.x.

Forde, T.L., Dennis, T.P.W., Aminu, O.R., Harvey, W.T., Hassim, A., Kiwelu, I., Medvecky, M., Mshanga, D., Van Heerden, H., Vogel, A., Zadoks, R.N., Mmbaga, B.T., Lembo, T. and Biek, R. (2022). Population genomics of Bacillus anthracis from an anthrax hyperendemic area reveals transmission processes across spatial scales and unexpected within-host diversity. Microbial Genomics, 8(2), 000759. doi:10.1099/mgen.0.000759.

Giani, T., Sennati, S., Antonelli, A., Di Pilato, V., di Maggio, T., Mantella, A., Niccolai, C., Spinicci, M., Monasterio, J., Castellanos, P., Martinez, M., Contreras, F., Balderrama Villaroel, D., Damiani, E., Maury, S., Rocabado, R., Pallecchi, L., Bartoloni, A. and Rossolini, G.M. (2018). High prevalence of carriage of mcr-1-positive enteric bacteria among healthy children from rural communities in the Chaco region, Bolivia, September to October 2016. Eurosurveillance, 23(45). doi:10.2807/1560-7917.es.2018.23.45.1800115.

Gilchrist, C. L., and Chooi, Y. H. (2021). Clinker & clustermap. js: Automatic generation of gene cluster comparison figures. Bioinformatics, 37(16), 2473–2475. doi: 10.1093/bioinformatics/btab007.

Gutiérrez, C., Zenis, J., Legarraga, P., Cabrera-Pardo, J.R., García, P., Bello-Toledo, H., Opazo-Capurro, A. and González-Rocha, G. (2019). Genetic analysis of the first mcr-1 positive Escherichia coli isolate collected from an outpatient in Chile. Brazilian Journal of Infectious Diseases, 23, 203–206. doi:10.1016/j.bjid.2019.05.008.

Hamel, M., Rolain, J.-M. and Baron, S.A. (2021). The History of Colistin Resistance Mechanisms in Bacteria: Progress and Challenges. Microorganisms, 9(2), 442. doi:10.3390/microorganisms9020442.

Holley, G., Beyter, D., Ingimundardottir, H., Møller, P.L., Kristmundsdottir, S., Eggertsson, H.P. and Halldorsson, B.V. (2021). Ratatosk: hybrid error correction of long reads enables accurate variant calling and assembly. Genome Biology, 22(1). doi:10.1186/s13059-020-02244-4.

Hong, Y.-K. and Ko, K.S. (2019). PmrAB and PhoPQ Variants in Colistin-Resistant Enterobacter spp. Isolates in Korea. Current Microbiology, 76(5), 644–649. doi:10.1007/s00284-019-01672-1.

Javed, H., Saleem, S., Zafar, A., Ghafoor, A., Shahzad, A.B., Ejaz, H., Junaid, K. and Jahan, S. (2020). Emergence of plasmid-mediated mcr genes from Gram-negative bacteria at the human-animal interface. Gut Pathogens, 12(1). doi:10.1186/s13099-020-00392-3.

Katip, W., Yoodee, J., Uitrakul, S. and Oberdorfer, P. (2021). Efficacy of loading dose colistin versus carbapenems for treatment of extended spectrum beta lactamase producing Enterobacteriaceae. Scientific Reports, 11(1). doi:10.1038/s41598-020-78098-4.

Kieffer, N., Royer, G., Decousser, J.-W., Bourrel, A.-S., Palmieri, M., Ortiz De La Rosa, J.-M., Jacquier, H., Denamur, E., Nordmann, P. and Poirel, L. (2019). mcr-9, an Inducible Gene Encoding an Acquired Phosphoethanolamine Transferase in Escherichia coli, and Its Origin. Antimicrobial Agents and Chemotherapy, 63(9). doi:10.1128/aac.00965-19.

Krutova, M., Kalova, A., Nycova, E., Gelbicova, T., Karpiskova, R., Smelikova, E., Nyc, O., Drevinek, P. and Tkadlec, J. (2021). The colonisation of Czech travellers and expatriates living in the Czech Republic by colistin-resistant Enterobacteriaceae and whole genome characterisation of E. coli isolates harbouring the mcr-1 genes on a plasmid or chromosome: A cross-sectional study. Travel Medicine and Infectious Disease, 39, p.101914. doi:10.1016/j.tmaid.2020.101914.

Larsen, M.V., Cosentino, S., Rasmussen, S., Friis, C., Hasman, H., Marvig, R.L., Jelsbak, L., Sicheritz-Ponten, T., Ussery, D.W., Aarestrup, F.M. and Lund, O. (2012). Multilocus Sequence Typing of Total-Genome-Sequenced Bacteria. Journal of Clinical Microbiology, 50(4), 1355–1361. doi:10.1128/jcm.06094-11.

Letunic, I. and Bork, P. (2021). Interactive Tree Of Life (iTOL) v5: an online tool for phylogenetic tree display and annotation. Nucleic Acids Research, 49(W1), W293–W296. doi:10.1093/nar/gkab301.

Li, R., Xie, M., Zhang, J., Yang, Z., Liu, L., Liu, X., Zheng, Z., Chan, E.W.-C. and Chen, S. (2016). Genetic characterization of mcr-1 -bearing plasmids to depict molecular mechanisms underlying dissemination of the colistin resistance determinant. Journal of Antimicrobial Chemotherapy, 72(2), 393–401. doi:10.1093/jac/dkw411.

Li, Y., Dai, X., Zeng, J., Gao, Y., Zhang, Z. and Zhang, L. (2020). Characterization of the global distribution and diversified plasmid reservoirs of the colistin resistance gene mcr-9. Scientific Reports, 10(1). doi:10.1038/s41598-020-65106-w.

Liao, W., Cui, Y., Quan, J., Zhao, D., Han, X., Shi, Q., Wang, Q., Jiang, Y., Du, X., Li, X. and Yu, Y. (2022). High prevalence of colistin resistance and mcr-9/10 genes in Enterobacter spp. in a tertiary hospital over a decade. International Journal of Antimicrobial Agents, 59(5), 106573. doi:10.1016/j.ijantimicag.2022.106573.

Lin, Y., Yuan, J., Kolmogorov, M., Shen, M.W., Chaisson, M. and Pevzner, P.A. (2016). Assembly of long error-prone reads using de Bruijn graphs. Proceedings of the National Academy of Sciences, 113(52), E8396–E8405. doi:10.1073/pnas.1604560113.

Ling, Z., Yin, W., Shen, Z., Wang, Y., Shen, J. and Walsh, T.R. (2020). Epidemiology of mobile colistin resistance genes mcr-1 to mcr-9. Journal of Antimicrobial Chemotherapy, 75(11), 3087– 3095. doi:10.1093/jac/dkaa205.

Loman, N.J. and Quinlan, A.R. (2014). Poretools: a toolkit for analyzing nanopore sequence data. Bioinformatics, 30(23), 3399–3401. doi:10.1093/bioinformatics/btu555.

Luo, Q., Wang, Y. and Xiao, Y. (2020). Prevalence and transmission of mobilized colistin resistance (mcr) gene in bacteria common to animals and humans. Biosafety and Health, 2(2), 71–78.doi:10.1016/j.bsheal.2020.05.001.

Marchetti, V.M., Bitar, I., Sarti, M., Fogato, E., Scaltriti, E., Bracchi, C., Hrabak, J., Pongolini, S. and Migliavacca, R. (2021). Genomic Characterization of VIM and MCR Co-Producers: The First Two Clinical Cases, in Italy. Diagnostics, 11(1), 79. doi:10.3390/diagnostics11010079.

Meier-Kolthoff, J.P., Auch, A.F., Klenk, H.-P. and Göker, M. (2013). Genome sequence-based species delimitation with confidence intervals and improved distance functions. BMC Bioinformatics, 14(1), 60. doi:10.1186/1471-2105-14-60.

Migura-Garcia, L., González-López, J.J., Martinez-Urtaza, J., Aguirre Sánchez, J.R., Moreno-Mingorance, A., Perez de Rozas, A., Höfle, U., Ramiro, Y. and Gonzalez-Escalona, N. (2020). mcr-Colistin Resistance Genes Mobilized by IncX4, IncHI2, and IncI2 Plasmids in Escherichia coli of Pigs and White Stork in Spain. Frontiers in Microbiology, 10. doi:10.3389/fmicb.2019.03072.

Quiroga, C., Nastro, M. and Di Conza, J. (2019). Current scenario of plasmid-mediated colistin resistance in Latin America. Revista Argentina de Microbiología, 51(1), 93–100. doi:10.1016/j.ram.2018.05.001.

Page, A.J., Cummins, C.A., Hunt, M., Wong, V.K., Reuter, S., Holden, M.T.G., Fookes, M., Falush, D., Keane, J.A. and Parkhill, J. (2015). Roary: rapid large-scale prokaryote pan genome analysis. Bioinformatics, 31(22), 3691–3693. doi:10.1093/bioinformatics/btv421.

Papagiannitsis, C.C., Študentová, V., Izdebski, R., Oikonomou, O., Pfeifer, Y., Petinaki, E. and Hrabák, J. (2015). Matrix-Assisted Laser Desorption Ionization–Time of Flight Mass Spectrometry Meropenem Hydrolysis Assay with NH 4 HCO 3, a Reliable Tool for Direct Detection of Carbapenemase Activity. Journal of Clinical Microbiology, 53(5), 1731–1735. doi:10.1128/jcm.03094-14.

Rebelo, A.R., Bortolaia, V., Kjeldgaard, J.S., Pedersen, S.K., Leekitcharoenphon, P., Hansen, I.M., Guerra, B., Malorny, B., Borowiak, M., Hammerl, J.A., Battisti, A., Franco, A., Alba, P., Perrin-Guyomard, A., Granier, S.A., De Frutos Escobar, C., Malhotra-Kumar, S., Villa, L., Carattoli, A. and Hendriksen, R.S. (2018). Multiplex PCR for detection of plasmid-mediated colistin resistance determinants, mcr-1, mcr-2, mcr-3, mcr-4 and mcr-5 for surveillance purposes. Euro surveillance, 23(6), 17–00672. doi:10.2807/1560-7917.ES.2018.23.6.17-00672.

Prim, N., Turbau, M., Rivera, A., Rodríguez-Navarro, J., Coll, P. and Mirelis, B. (2017). Prevalence of colistin resistance in clinical isolates of Enterobacteriaceae: A four-year cross-sectional study. Journal of Infection, 75(6), 493–498. doi:10.1016/j.jinf.2017.09.008.

Ruan, Z., Sun, Q., Jia, H., Huang, C., Zhou, W., Xie, X. and Zhang, J. (2019). Emergence of a ST2570 Klebsiella pneumoniae isolate carrying mcr-1 and blaCTX-M-14 recovered from a bloodstream infection in China. Clinical Microbiology and Infection, 25(7), 916–918. doi:10.1016/j.cmi.2019.02.005.

Skov, R.L. and Monnet, D.L. (2016). Plasmid-mediated colistin resistance (mcr-1 gene): three months later, the story unfolds. Eurosurveillance, 21(9). doi:10.2807/1560-7917.es.2016.21.9.30155.

Sun, J., Zeng, X., Li, X.-P., Liao, X.-P., Liu, Y.-H. and Lin, J. (2017). Plasmid-mediated colistin resistance in animals: current status and future directions. Animal Health Research Reviews, 18(2), 136–152. doi:10.1017/S1466252317000111.

Tijet, N., Faccone, D., Rapoport, M., Seah, C., Pasterán, F., Ceriana, P., Albornoz, E., Corso, A., Petroni, A. and Melano, R.G. (2017). Molecular characteristics of mcr-1-carrying plasmids and new mcr-1 variant recovered from polyclonal clinical Escherichia coli from Argentina and Canada. PLoS One, 12(7), e0180347. doi:10.1371/journal.pone.0180347.

Tkadlec, J., Kalova, A., Brajerova, M., Gelbicova, T., Karpiskova, R., Smelikova, E., Nyc, O., Drevinek, P. and Krutova, M. (2021). The Intestinal Carriage of Plasmid-Mediated Colistin-Resistant Enterobacteriaceae in Tertiary Care Settings. Antibiotics, 10(3), 258. doi:10.3390/antibiotics10030258.

Tyson, G.H., Li, C., Hsu, C.-H., Ayers, S., Borenstein, S., Mukherjee, S., Tran, T.-T., McDermott, P.F. and Zhao, S. (2020). The mcr-9 Gene of Salmonella and Escherichia coli Is Not Associated with Colistin Resistance in the United States. Antimicrobial Agents and Chemotherapy, 64(8). doi:10.1128/aac.00573-20.

Viñes, J., Cuscó, A., Napp, S., Alvarez, J., Saez-Llorente, J.L., Rosàs-Rodoreda, M., Francino, O. and Migura-Garcia, L. (2021). Transmission of Similar mcr-1 Carrying Plasmids among Different Escherichia coli Lineages Isolated from Livestock and the Farmer. Antibiotics, 10(3), 313. doi:10.3390/antibiotics10030313.

Walker, B.J., Abeel, T., Shea, T., Priest, M., Abouelliel, A., Sakthikumar, S., Cuomo, C.A., Zeng, Q., Wortman, J., Young, S.K. and Earl, A.M. (2014). Pilon: An Integrated Tool for Comprehensive Microbial Variant Detection and Genome Assembly Improvement. PLoS ONE, 9(11), e112963. doi:10.1371/journal.pone.0112963.

Wand, M.E. and Sutton, J.M. (2020). Mutations in the two component regulator systems PmrAB and PhoPQ give rise to increased colistin resistance in Citrobacter and Enterobacter spp. Journal of Medical Microbiology, 69(4), 521–529. doi:10.1099/jmm.0.001173.

Wang, X., Wang, Y., Zhou, Y., Li, J., Yin, W., Wang, S., Zhang, S., Shen, J., Shen, Z. and Wang, Y. (2018). Emergence of a novel mobile colistin resistance gene, mcr-8, in NDM-producing Klebsiella pneumoniae. Emerging Microbes & Infections, 7(1), 1–9. doi:10.1038/s41426-018-0124-z.

Wang, C., Feng, Y., Liu, L., Wei, L., Kang, M. and Zong, Z. (2020). Identification of novel mobile colistin resistance gene mcr-10. Emerging Microbes & Infections, 9(1), 508–516. doi:10.1080/22221751.2020.1732231.

Wick, R.R., Judd, L.M., Gorrie, C.L. and Holt, K.E. (2017). Unicycler: Resolving bacterial genome assemblies from short and long sequencing reads. PLoS Computational Biology, 13(6), e1005595. doi:10.1371/journal.pcbi.1005595.

Xavier, B.B., Lammens, C., Butaye, P., Goossens, H. and Malhotra-Kumar, S. (2016). Complete sequence of an IncFII plasmid harbouring the colistin resistance gene mcr-1 isolated from Belgian pig farms. Journal of Antimicrobial Chemotherapy, 71(8), 2342–2344. doi:10.1093/jac/dkw191.

Yoon, S. H., Ha, S. M., Kwon, S., Lim, J., Kim, Y., Seo, H. and Chun, J. (2017). Introducing EzBioCloud: a taxonomically united database of 16S rRNA gene sequences and whole-genome assemblies. International journal of systematic and evolutionary microbiology, 67(5), 1613. doi: 10.1007/s10482-017-0844-4.

Yilmaz, G.R., Dizbay, M., Guven, T., Pullukcu, H., Tasbakan, M., Guzel, O.T., Tekce, Y.T., Ozden, M., Turhan, O., Guner, R., Cag, Y., Bozkurt, F., Karadag, F.Y., Kartal, E.D., Gozel, G., Bulut, C., Erdinc, S., Keske, S., Acikgoz, Z.C. and Tasyaran, M.A. (2016). Risk factors for infection with colistin-resistant gram-negative microorganisms: a multicenter study. Annals of Saudi Medicine, 36(3), 216–222. doi:10.5144/0256-4947.2016.216.

Zankari, E., Hasman, H., Cosentino, S., Vestergaard, M., Rasmussen, S., Lund, O., Aarestrup, F.M. and Larsen, M.V. (2012). Identification of acquired antimicrobial resistance genes. Journal of Antimicrobial Chemotherapy, 67(11), 2640–2644. doi:10.1093/jac/dks261.

Zankari, E., Allesøe, R., Joensen, K.G., Cavaco, L.M., Lund, O. and Aarestrup, F.M. (2017). PointFinder: a novel web tool for WGS-based detection of antimicrobial resistance associated with chromosomal point mutations in bacterial pathogens. Journal of Antimicrobial Chemotherapy, 72(10), 2764–2768. doi:10.1093/jac/dkx217.

Zelendova, M., Papagiannitsis, C.C., Valcek, A., Medvecky, M., Bitar, I., Hrabak, J., Gelbicova, T., Barakova, A., Kutilova, I., Karpiskova, R. and Dolejska, M. (2021). Characterization of the Complete Nucleotide Sequences of mcr-1-Encoding Plasmids From Enterobacterales Isolates in Retailed Raw Meat Products From the Czech Republic. Frontiers in Microbiology, 11. doi:10.3389/fmicb.2020.604067.

Zhang, Z., Tian, X. and Shi, C. (2022). Global Spread of MCR-Producing Salmonella enterica Isolates. Antibiotics (Basel, Switzerland), 11(8), 998. doi:10.3390/antibiotics11080998.

Zingali, T., Chapman, T.A., Webster, J., Roy Chowdhury, P. and Djordjevic, S.P. (2020). Genomic Characterisation of a Multiple Drug Resistant IncHI2 ST4 Plasmid in Escherichia coli ST744 in Australia. Microorganisms, 8(6), 896. doi:10.3390/microorganisms8060896.

Zhu, W., Lawsin, A., Lindsey, R.L., Batra, D., Knipe, K., Yoo, B.B., Perry, K.A., Rowe, L.A., Lonsway, D., Walters, M.S., Rasheed, J.K. and Halpin, A.L. (2019). Conjugal Transfer, Whole-Genome Sequencing, and Plasmid Analysis of Four mcr-1 -Bearing Isolates from U.S. Patients. Antimicrobial Agents and Chemotherapy, 63(4). doi:10.1128/aac.02417-18.

Zurfluh, K., Nüesch-Inderbinen, M., Klumpp, J., Poirel, L., Nordmann, P. and Stephan, R. (2017). Key features of mcr-1-bearing plasmids from Escherichia coli isolated from humans and food. Antimicrobial Resistance & Infection Control, 6(1). doi:10.1186/s13756-017-0250-8.

