## Supplementary figures and images for "Plasmid-mediated colistin resistance among human clinical *Enterobacterales* isolates: National surveillance in the Czech Republic"

### Supplementary Figure 1

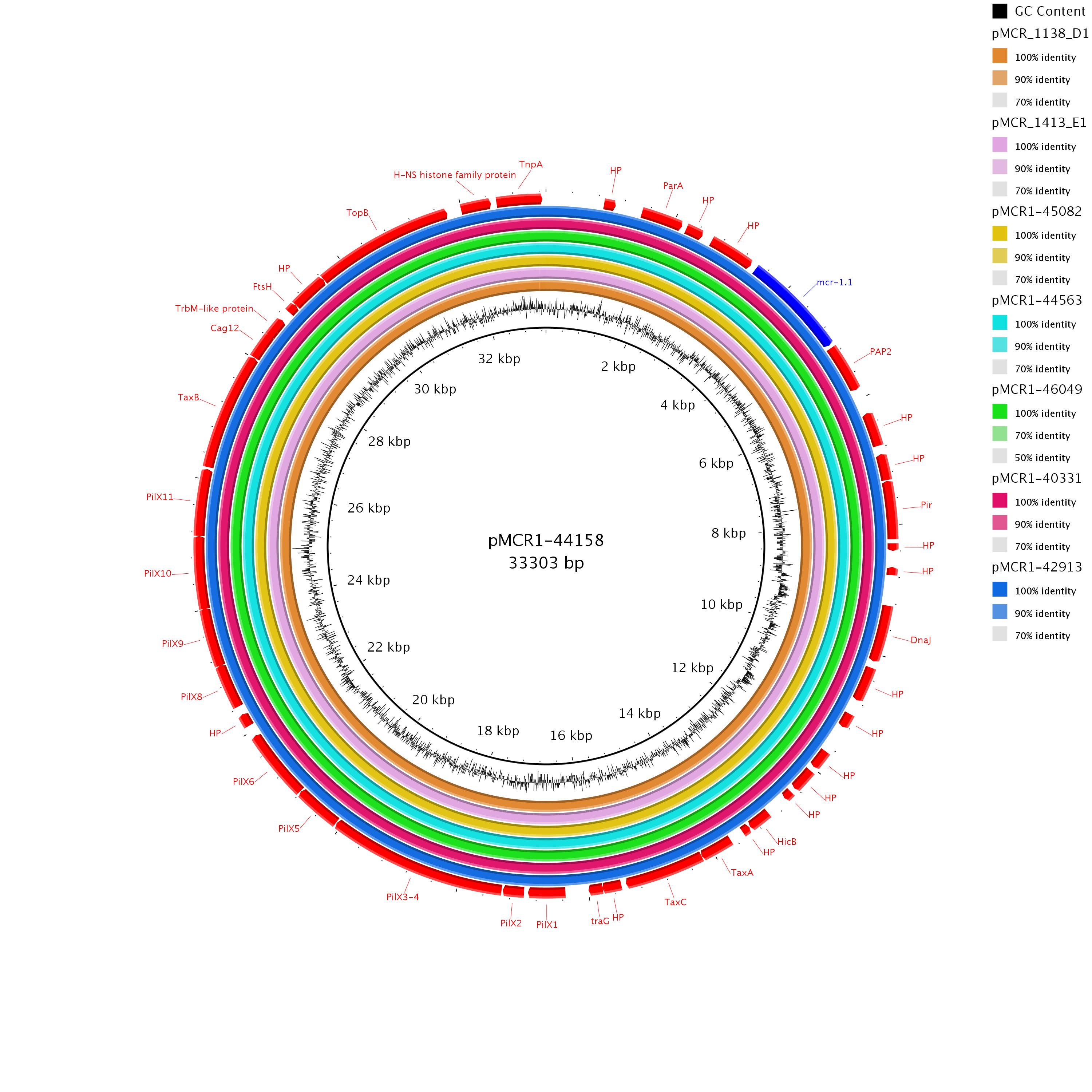

### Supplementary Figure 2

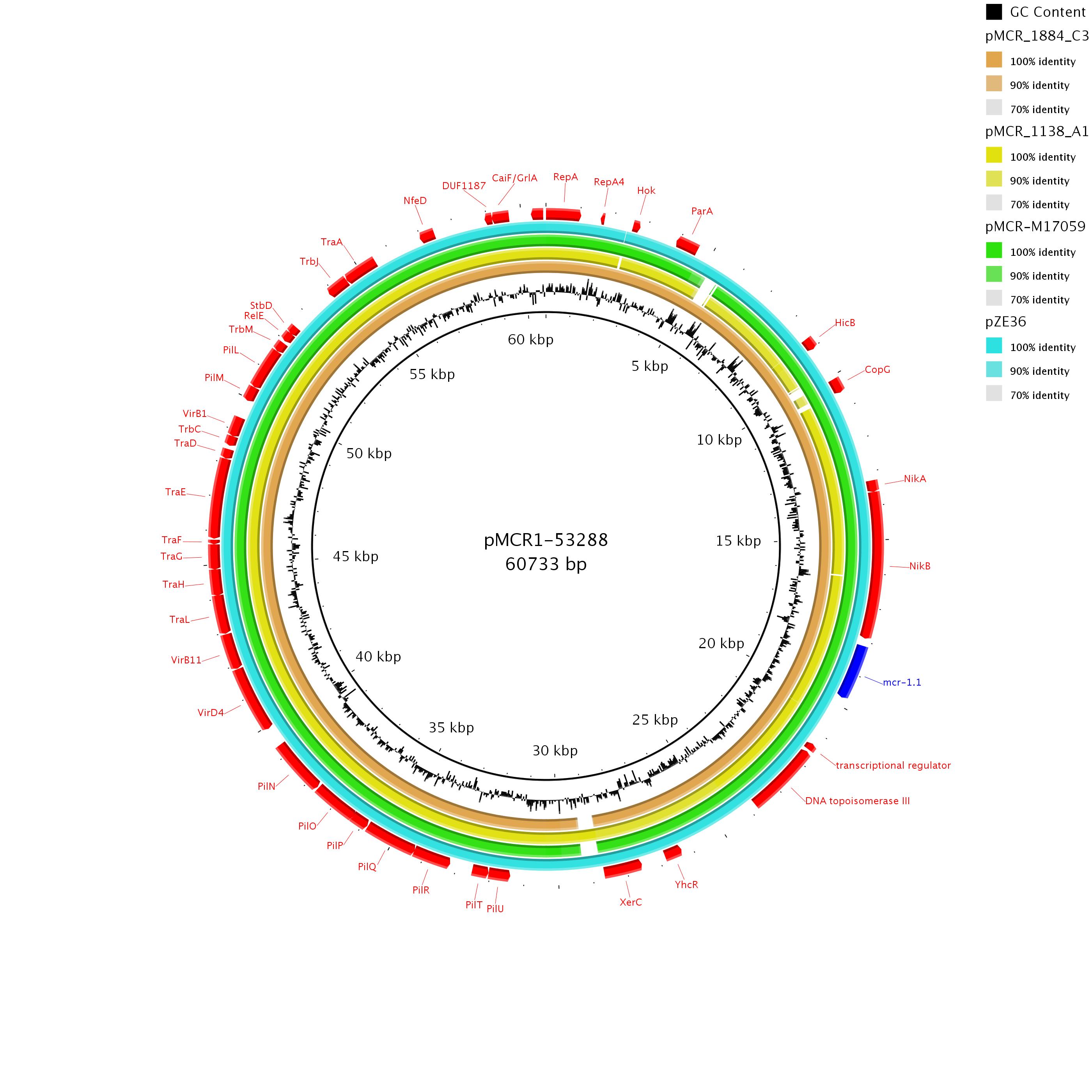

### Supplementary Figure 3

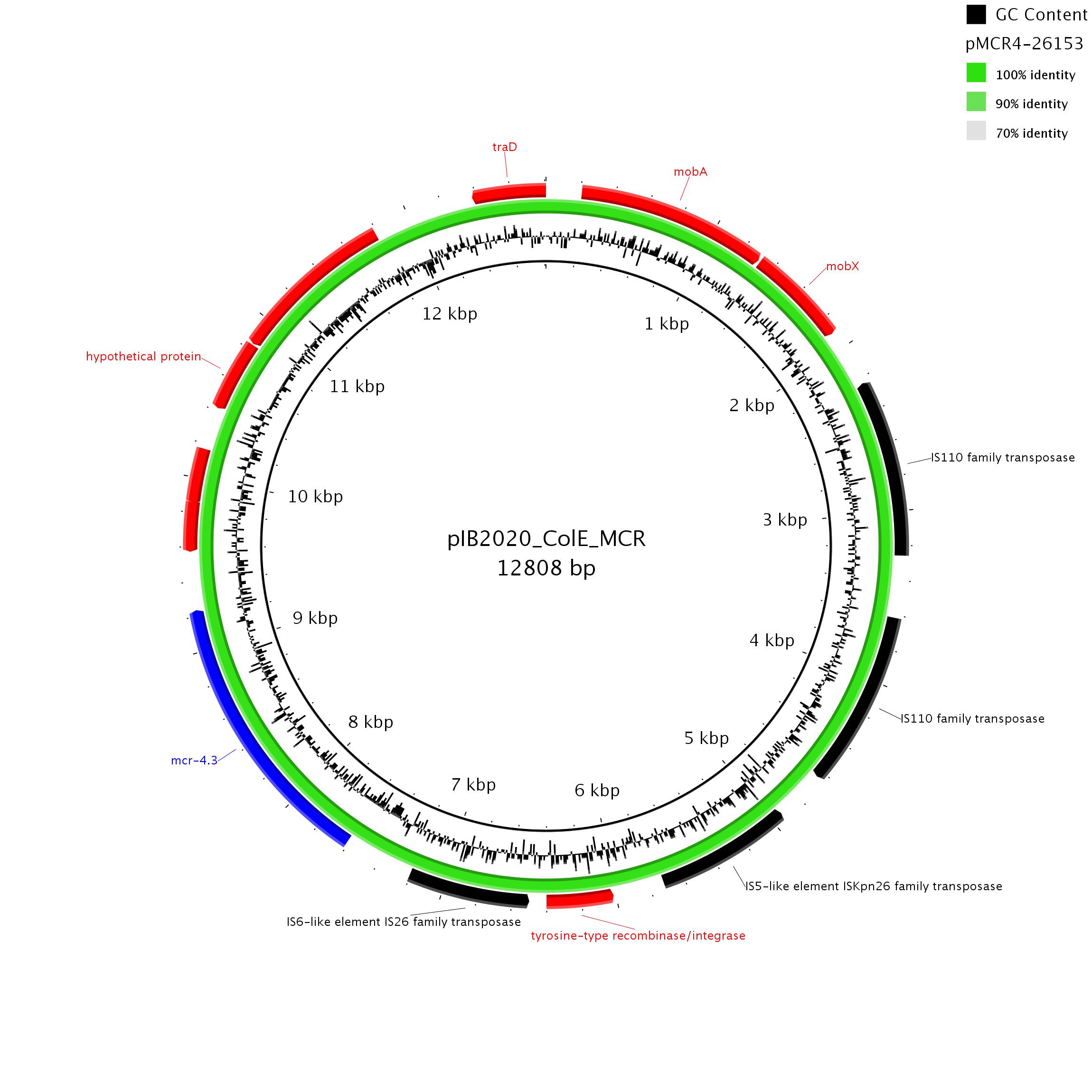

### Supplementary Figure 4

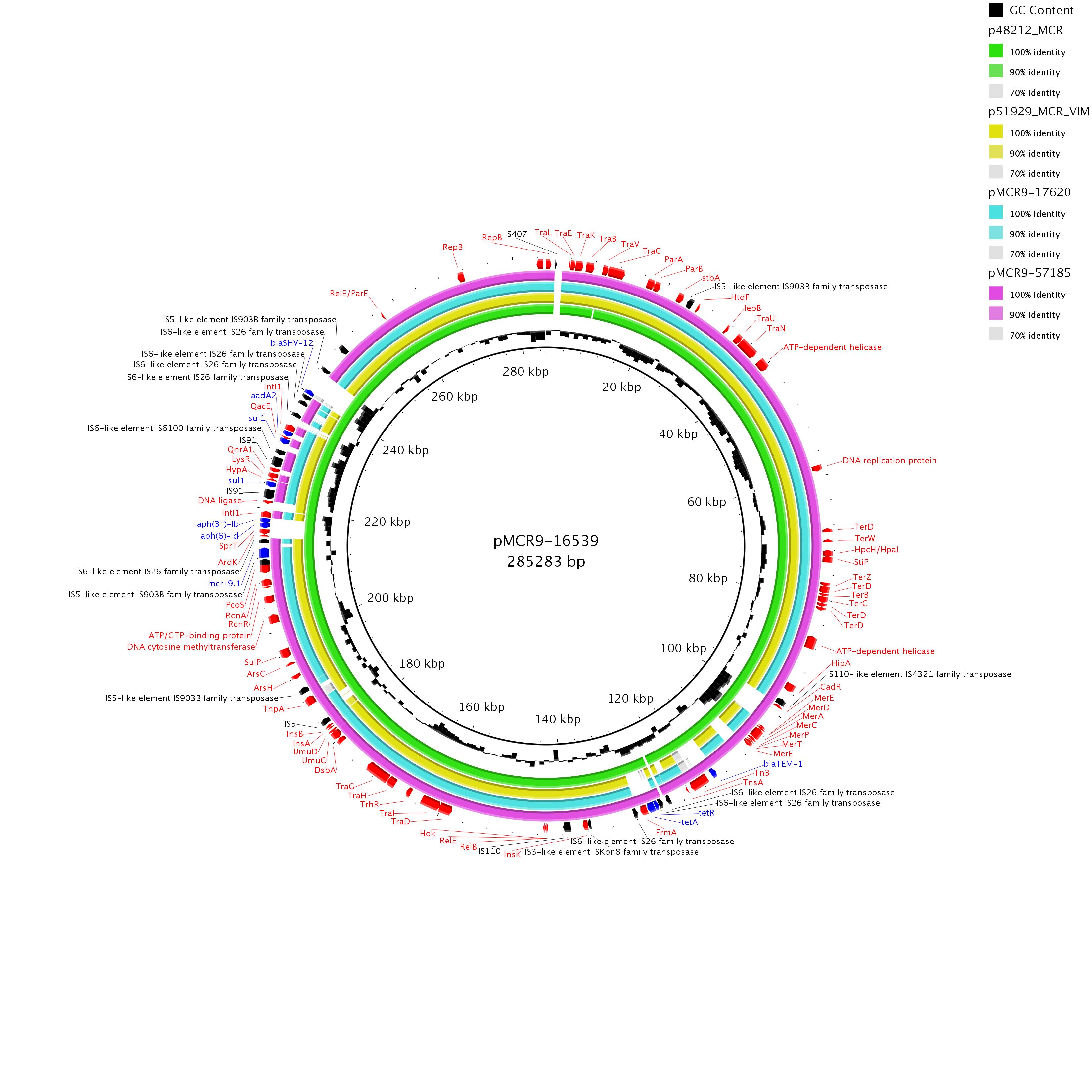

### Supplementary Figure 5

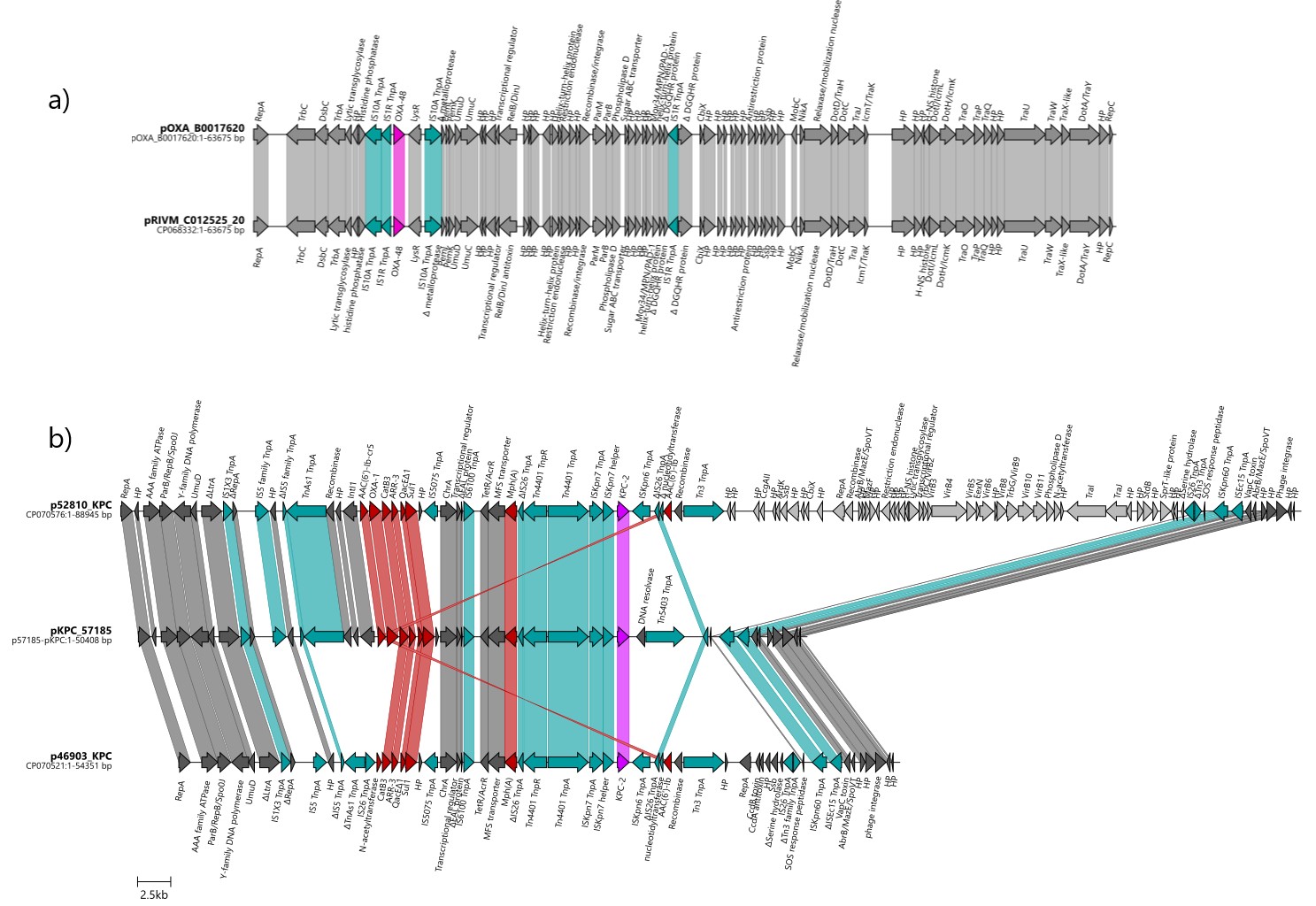
